# NMR and molecular dynamics demonstrate the RNA internal loop GAGU is dynamic and adjacent base pairs determine conformational preference

**DOI:** 10.1101/2025.04.14.648754

**Authors:** Olayinka Akinyemi, Scott D. Kennedy, James P. McSally, David H. Mathews

## Abstract

The conformational variability of RNA duplexes with the internal loop 5’GAGU/3’UGAG was investigated by a combination of nuclear magnetic resonance spectroscopy (NMR) and all-atom molecular dynamics (MD) simulations. A previous study showed that the CG-flanked internal loop in the sequence 5’GAC**GAGU**GUCA/3’ACUG**UGAG**CAG existed in a major conformation (conformation I) characterized by Uridines U7 and U7* bulging into solution, A5 and A5* stacking, and with G4-G6* and G6-G4* base pairs closing the loop on either end. It was also determined that a minor conformation existed with a set of closing G-U pairs with a bifurcated hydrogen bond and A-G non-canonical pairs with a single hydrogen bond (conformation II). A maximum hydrogen bonding structure with wobble G-U pairs and A-G pairs with two hydrogen bonds (conformation III) was not observed by NMR or predicted by molecular dynamics. In this work, alternative flanking base pairs A-U, U-A and G-C were studied by substituting the C-G base pair adjacent to the GAGU internal loop. NMR spectra demonstrated changes in conformational preference depending on the identity of the flanking pair. Clustering analyses of structures from MD simulations of C-G- and U-A-flanked duplexes showed transitions between minor conformations II and III with a greater fraction of the structures in conformation II, in spite of the simulations starting in conformation III. In addition, U-A-flanked simulations contained a substantial amount (25%) of structures in an intermediate state between conformations II and III. A-U- and G-C-flanked structures were all in a single cluster whose centroid structure was in conformation II. MD simulations showed a dominance of structure II over structure III, in agreement with NMR data for C-G closure, but in contrast with the NMR data for other closures. Simulations starting in conformation I did not transition to either of II and III, with all structures being in a single cluster for all flanking base pairs except for G-C-closure where 22% of the structures were in a state defined by a C3’endo sugar pucker for G4 and G4*. Bases G4 and G4* were also in syn orientation around the glycosidic bond for all flanking pairs while G6 and G6* transitioned between syn and anti orientations for C-G- and A-U-flanked simulations, consistent with NMR observations.

## INTRODUCTION

RNA serves numerous roles in cells, including catalysis, post-transcriptional gene regulation, and target recognition (1–5). To perform these roles, RNA adopts specific structures and these structures are dynamic, i.e. they sample multiple conformations at equilibrium. It is this dynamic nature that facilitates catalysis and interactions with ligands.

To better understand RNA structure and function, there is substantial effort to elucidate dynamics. Experimental methods can be used to understand dynamics, including single molecule fluorescence (6,7), NMR (8–10), and cryo-EM (11,12). Computationally, molecular dynamics (MD) simulations or quantum dynamics simulations are used to model dynamics from first principles (13,14). The experimental and computational methods work in tandem to provide a complete model of dynamics across timescales.

A model dynamic RNA structure is the internal loop formed by duplexes with the sequence GAGU_2_. NMR demonstrated that GAGU is dynamic and interconverts between a major conformation and a minor conformation, where the major conformation has a structure in which the Adenines stack, two G-G pairs form, and the Uridines flip out into solution (Fig. 1A, Conformation I) (15). This was unexpected because solution and crystal structures of similar 2×2 internal loops showed maximal non-canonical pairing of the nucleotides (16–20). The minor conformation was subsequently elucidated (Fig. 1A, Conformation II) (21). In that work, MD simulations showed a set of G-U pairs with a bifurcated hydrogen bond and A-G non-canonical pairs with a single hydrogen bond. The structure found by these simulations was supported by the NMR data as the GAGU minor conformation.

**Figure 1.**
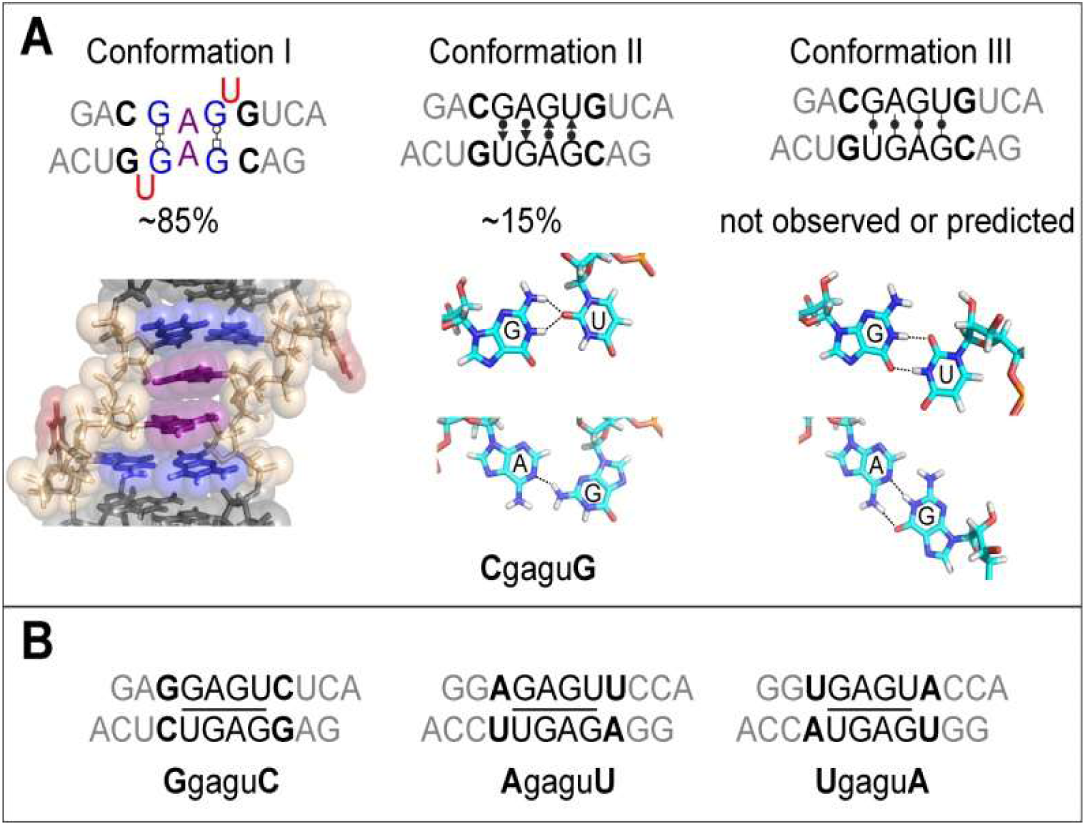
(A) Summary of conformations of (GA**C**<u>GAGU</u>**G**UCA)_2_ determined in earlier work (PDB ID 2LX1) (15). (B) Sequences and naming convention of three variations of GAGU studied in this work with adjacent base pairs other than CG (57).

Here we study three new variations of GAGU with alternative adjacent base pairs. The prior experiments and simulations were performed with C-G base pairs adjacent to the GAGU sequences (Fig. 1A). In this work, we substitute the C-G pairs with G-C, U-A, and A-U pairs (Fig. 1B) to improve our understanding of the relationships between sequence and structure. For brevity, the sequences are named XgaguY, where XY represents the adjacent base pair, i.e. CgaguG, GgaguC, AgaguU and UgaguA (Fig. 1). Interestingly, NMR demonstrates substantial changes in the conformational selection driven by the adjacent base pairs. Molecular dynamics simulations using the Amber force field (22–24), however, do not always correctly model the conformational preferences, suggesting a force field deficiency. The GAGU internal loop is therefore a good benchmark structure with which to test improvements in RNA force fields.

## MATERIALS AND METHODS

### NMR Samples, Data and Processing

Four RNA oligomers with sequences GA**C**<u>GAGU</u>**G**UCA, GA**G**<u>GAGU</u>**C**UCA, GG**A**<u>GAGU</u>**U**CCA, and GG**U**<u>GAGU</u>**A**CCA were obtained from Dharmacon, Inc. (now Horizon Discovery; Lafayette, CO), and prepared in aqueous buffer containing 70 mM sodium chloride, 30 mM sodium phosphate, and 0.1 mM EDTA at pH 6.1. RNA strand concentrations of the samples for NMR studies were 2.0 mM for CgaguG and GgaguC and 3.0 mM for AgaguU and UgaguA in 500 μl volume with 4% D_2_O. Samples at low concentration were made by 100-fold dilution with buffer. NMR spectra were acquired on a Varian (Agilent) Unity Inova spectrometer operating at 600 MHz for protons with a room temperature, triple resonance, three axis gradient probe. One-dimensional spectra and two-dimensional NOESY spectra using a WATERGATE pulse with flipback for water suppression were acquired at 1, 10, and 20 °C (25,26). NOESY spectra with various mixing times provided most of the needed assignment and structure information with assist or confirmation from 2D TOCSY and 1H-13C HSQC. Reference to previous experiments and analyses (15,17,21) of CgaguG provided guidance assigning the conformations in all four sequences. Spectra from 1D NMR were analyzed on the spectrometer (VNMR version 6.1C). Spectra from 2D NMR were processed, assigned, and integrated with NMRPipe (27) and NMRFAM-SPARKY (28).

Relative populations of conformation I versus population of the alternative conformation (population of conformation II, III, or a mixture of II and III; Fig. 2) were determined by comparing 1D imino peak areas and 2D NOESY cross-peak volumes associated with the Watson-Crick-Franklin base pairs adjacent to the terminal G1-C10* pairs (A2-U9* pair for CgaguG and GgaguC; G2-C9* pair for UgaguA and AgaguU). These pairs were chosen for their lack of dynamics and sufficient spectral resolution between conformation I and conformation II/III.

**Figure 2.**
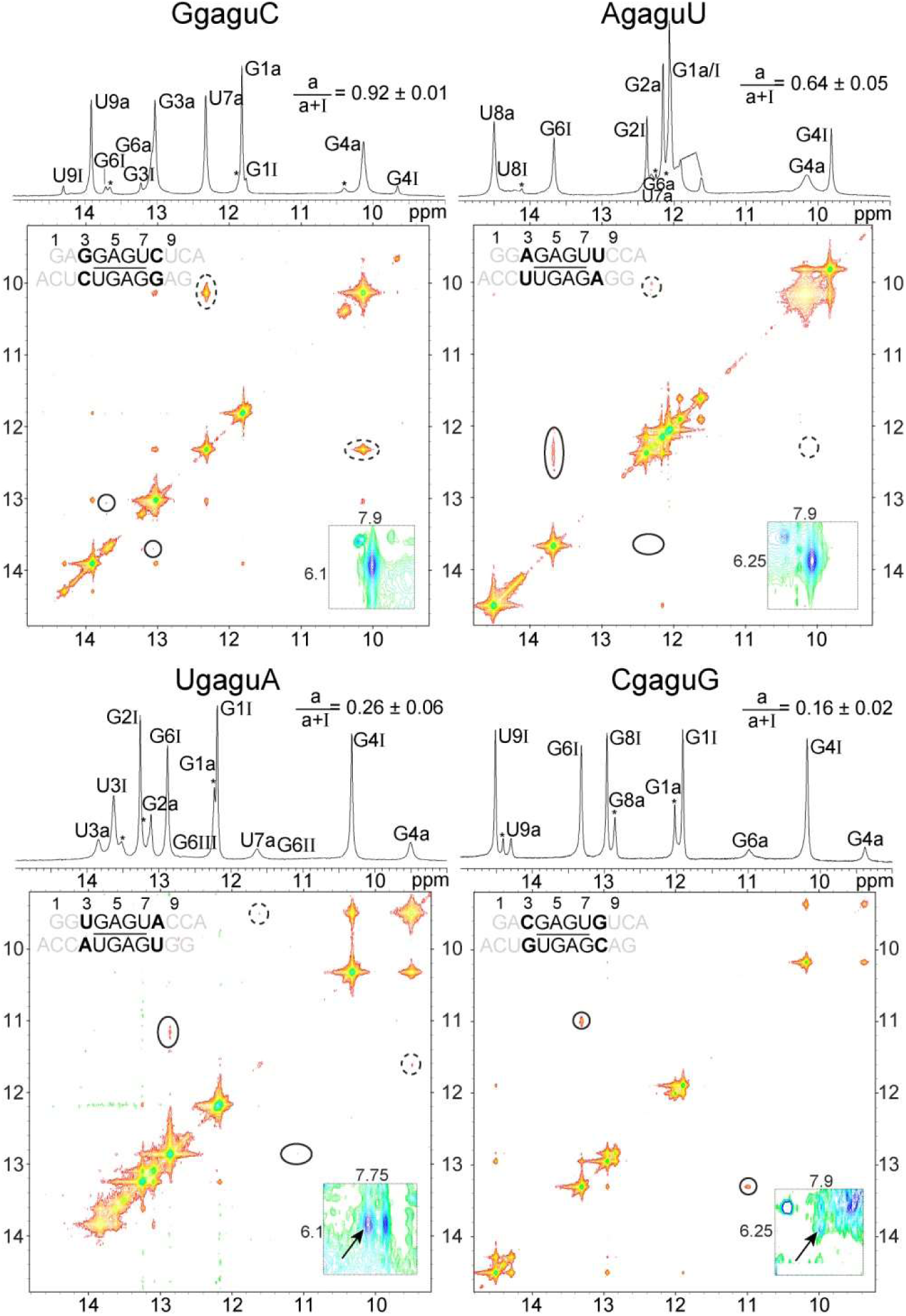
NMR spectra of imino proton region of four RNA duplexes at 1 °C. Peaks are identified as belonging to either conformation I (I) or an alternative conformation (a) which is either conformation II, conformation III, or a mixture of conformation II and III (Fig. 1). Peaks marked with an asterisk are due to a single-strand (hairpin) population. The population of conformation I relative to the alternative conformation is indicated above each spectrum. 2D NOESY spectra are shown below each 1D spectrum. Circled cross peaks indicate exchange between alternative conformation and conformation I (solid line) or U7H3-G4*H1 in GU wobble of conformation III (dashed line). Insets show A5H2-G6*amino cross-peaks (arrows) with contour intensities scaled to C10 amino-amino cross-peak intensities that are independent of conformation, and inversely scaled by the population of the alternative conformation.

Linewidths were measured (in Hz) using the 2D peak fitting function in Sparky. The strongest available NOESY cross-peaks of aromatic or G6 imino protons were chosen for fitting peaks. Linewidth averages and uncertainties were determined from four spectra of each duplex.

Volumes of G4H8-H1’ and G6H8-H1’ NOESY cross-peaks of each duplex were measured in two spectra, 75 and 50 ms mixing time, by the 2D fitting tool in NMRFAM-SPARKY. The average of the highest and lowest volumes from four data range selections were used to obtain a ratio of the G4 and G6 volumes.

### NMR-guided modeling of GgaguC

Structure models of GgaguC were generated by simulated annealing with AMBER 2016 (29) using distance and dihedral angle restraints generated from NMR data. The calculation started with a model derived from the major conformation of CgaguG (conformation I). A generalized Born implicit solvent was used with a salt concentration of 0.1 M NaCl (30). The temperature was started at 3000 K, then reduced to 0 K over 100 ps of simulation. Full NOE and dihedral scale factors were 30 kcal/mol.Å^2^ and 30 kcal/mol.rad^2^, respectively. Dihedral angles alpha and zeta were not restrained within the GAGU internal loop. This procedure was repeated 600 times with different velocity seeds. Structures with the 30 lowest restraint energies and no distance violations greater than 0.1 Å were selected for the final ensemble deposited in the RCSB Protein Data Bank with ID code 9BXE.

### Preparation of MD models

A total of eight duplex models, composed of the four sequences containing the GAGU internal loop with differing adjacent base pairs and starting in either conformation I or conformation III (Fig. 1), were simulated using all-atom MD simulation with the Amber force field (22–24). Each model was simulated with four independent simulations for 1 μs each.

Starting models for conformation I of GgaguC, AgaguU and UgaguA were generated as derivatives of conformation I of CgaguG (PDB ID 2LX1) (15) by correspondingly swapping out the CG sequence for the XY sequence through sequence mutation in Chimera (31). Starting models for conformation III were derived from the structure of the GAGC internal loop (PDB ID 2KY0) (17). The cytosine was mutated to a uracil to produce a conformation III model of CgaguG followed by mutation of the CG sequence to the corresponding XY sequence. The starting structures were neutralized with K^+^ counterions (32) and then solvated in OPC water (33) in a truncated octahedron box with periodic boundary condition and a 10 Å margin of buffer. Additional K^+^ and Cl^−^ ions were added to bring the molarity to 0.1 M (34). A separate set of starting structures were prepared in 1 M K^+^ and Cl^−^ in excess of neutrality as a control to understand possible conformational changes due to a change in salt concentration.

### Simulation protocol

The four replicates of each sequence were simulated by all-atom molecular dynamics using the Amber GPU code (35) and the standard AMBER force field (RNA.OL3) (22–24). The solvated starting structures were energy-minimized to bring the structures to the nearest local potential energy minimum. The minimization was a two-stage process, with the first stage carried out by placing a weak harmonic restraint (1.0 kcal·mol^−1^·Å^−2^) on the heavy atoms while allowing water and lighter atoms to minimize, followed by the removal of the restraint in the second minimization stage. Both minimization steps used 500 steps of steepest descent followed by 500 steps of conjugate gradient minimization. Heating was done at NVT to raise the temperature of the system to 10 °C over a period of 200 ps. The system was equilibrated for 1 ns followed by a 1 μs production molecular dynamics, with both processes carried out at constant temperature and pressure. Temperature regulation was implemented using a Langevin thermostat (36) with a collision frequency of 2 ps^−1^ while a Monte-Carlo barostat (37) with a 100- step attempt frequency was used to regulate the pressure. Particle mesh Ewald was used for periodic boundary conditions, and the direct space sum was limited to a cutoff of 8.0 Å. Hydrogen motion was constrained with SHAKE (38), allowing for a time step of 2 fs. Snapshots of the trajectory were recorded every 50,000 time steps, corresponding to 200 ps intervals.

### MD analysis methods

Postprocessing of trajectories was carried out in CPPTRAJ, which is contained in the Amber suite (39). Trajectories were centered and reimaged into the primary simulation box to prevent artifacts associated with the use of periodic boundary conditions. To investigate convergence and possible conformational changes, root mean square deviation (RMSD) relative to the energy-minimized structure was computed and plotted. K-Means clustering was used to group the structures into clusters of similar conformations. The atomic coordinate dataset was subjected to dimensionality reduction through principal component analysis (PCA), retaining the first ten principal components, which accounted for at least 80% of the total variance in the data. This was necessary to aid the convergence of the K-Means algorithm, with the optimal number of clusters determined using the elbow method (40,41). Specific dihedral angles and distances were computed for the centroid structure of GgaguC to detect conformations of interest such as crankshafted backbones (γ and α) that were detected in 2D NOE of the sequence (Table S4), and *syn* bases (χ).

## RESULTS

### NMR Experiments

NMR spectra of each sequence demonstrated the presence of multiple duplex conformations (Fig. 2). In addition, a small amount (< 10% at 1 °C) of single stranded hairpin structure was present in each sample, where the hairpins were identified in NMR spectra as peaks that became more intense relative to duplex signals following ∼100 fold dilution of the concentrated samples (Fig. S1; see Hammond et al. (17) for CgaguG dilution).

As duplexes, all four sequences formed the same 4×4 internal loop, 5’GAGU/3’UGAG, with different Watson-Crick-Franklin closing pairs. The internal loop of each sequence exists in two conformations that are in moderately slow exchange (5-200 ms) as evidenced in 2D NOESY spectra by an exchange cross-peak between peaks of each conformation (Fig. 2). One of the two slowly exchanging conformations is the conformation originally identified by Hammond et al. (17) with the structure determined by Kennedy et al. (15). Here we refer to this as conformation I as shown in Fig. 1.

Typical NOEs that identify conformation I include strong G4H8-H1’ and G6H8-H1’ due to the *syn* conformation of G4 and G6, strong G4H8-G6*H1 due to G4/G6* Watson- Crick/Hoogsteen pairs (* represents the opposite strand), and G4H1-A5*H2. Downfield shifts of U7H6 and U7H5 due to a flipped out U7 are also characteristic. The similarity of the conformation I structures observed in the four sequences is demonstrated by the highly correlated chemical shifts of atoms near the center of the loop, away from the influence of the differing closing pairs (Table 1). Based on an analysis of the volume of G4H8-H1’ and G6H8-H1’ cross-peaks, G6 χ exhibits a preference relative to G4 χ for a non-syn orientation. The ratio of the G6H8-H1’ volume to G4H8-H1’ volume is 0.90±0.02 for AgaguU, 0.83±0.01 for CgaguG, and 0.60±0.04 for UgaguA, indicating a lower proportion of syn G6 χ relative to syn G4 χ for these duplexes (Table S2). The low population of conformation I prevents determination of this ratio in GgaguC.

**Table 1.** Chemical shifts of protons within the GAGU internal loop that are not adjacent to the closing Watson-Crick-Franklin base pairs at 1, 10, and 20 °C. Appropriate protons for conformation I and the alternative conformations (II / III) are reported separately. Standard deviation of chemical shifts is included to reflect the similarity or dissimilarity of a given conformation between the four sequences (small deviation indicates high structural similarity). With the assumption that conformation II and III are the only alternative conformations present, percent of conformation II or III are calculated from average of A5H8, A5H1’, G6H8, and G6H1 and variance of those averages assuming that CgaguG represents 100% conformation II and GgaguC represents 100% conformation III. A complete list of all assigned atom shifts at 1 °C can be found in Table S1.

| 1 °C |  | A5H8 | A5H1' | A5H2 |  |  |  |  |
| --- | --- | --- | --- | --- | --- | --- | --- | --- |
| Conformation I | GgaguC | 7.55 | 5.7 | 7.69 |  |  |  |  |
|  | AgaguU | 7.57 | 5.65 | 7.68 |  |  |  |  |
|  | UgaguA | 7.6 | 5.69 | 7.69 |  |  |  |  |
|  | CgaguG | 7.58 | 5.69 | 7.67 |  |  |  |  |
| std dev |  | 0.021 | 0.022 | 0.010 |  |  |  |  |
|  |  |  |  |  | G6H8 | G6H1' | G6H1 | % II / III |
| Conformation II/III | GgaguC | 8.16 | 6.01 | 7.89 | 7.07 | 5.48 | 13.06 | 0 / 100 |
|  | AgaguU | 8 | 5.9 | 7.88 | 7.22 | 5.42 | 12.35 | 33 / 67 ± 3 |
|  | UgaguA | 8.08 | 5.89 | 7.75 | 7.16 | 5.4 | 12.48 | 24 / 76 ± 7 |
|  | CgaguG | 7.69 | 5.63 | 7.91 | 7.51 | 5.39 | 11 | 100 / 0 |
| std dev |  | 0.206 | 0.161 | 0.073 | 0.190 | 0.040 | 0.871 |  |

| 10 °C |  | A5H8 | A5H1' | A5H2 |  |  |  |  |
| --- | --- | --- | --- | --- | --- | --- | --- | --- |
| Conformation I | GgaguC | 7.56 | 5.7 | 7.7 |  |  |  |  |
|  | AgaguU | 7.56 | 5.65 | 7.69 |  |  |  |  |
|  | UgaguA | 7.6 | 5.68 | 7.69 |  |  |  |  |
|  | CgaguG | 7.59 | 5.69 | 7.69 |  |  |  |  |
| std dev |  | 0.021 | 0.022 | 0.005 |  |  |  |  |
|  |  |  |  |  | G6H8 | G6H1' | G6H1 | % II / III |
| Conformation II/III | GgaguC | 8.14 | 6.013 | 7.89 | 7.11 | 5.49 | 13.05 | 0 / 100 |
|  | AgaguU | 7.94 | 5.86 | 7.88 | 7.3 | 5.43 | 12.01 | 45 / 55 ± 4 |
|  | UgaguA | 8.03 | 5.85 | 7.75 | 7.22 | 5.42 | 12.37 | 31 / 69 ± 7 |
|  | CgaguG | 7.71 | 5.62 | 7.89 | 7.53 | 5.42 | 10.89 | 100 / 0 |
| std dev |  | 0.183 | 0.162 | 0.068 | 0.178 | 0.034 | 0.903 |  |

| 20 °C |  | A5H8 | A5H1' | A5H2 |  |  |  |  |
| --- | --- | --- | --- | --- | --- | --- | --- | --- |
| Conformation I | GgaguC | 7.57 | 5.72 | 7.72 |  |  |  |  |
|  | AgaguU | 7.57 | 5.66 | 7.7 |  |  |  |  |
|  | UgaguA |  |  |  |  |  |  |  |
|  | CgaguG | 7.56 | 5.68 |  |  |  |  |  |
| std dev |  | 0.006 | 0.031 | 0.014 |  |  |  |  |
|  |  |  |  |  | G6H8 | G6H1' | G6H1 | % II / III |
| Conformation II/III | GgaguC | 8.14 | 6.02 | 7.89 | 7.16 | 5.53 | 12.9 | 0 / 100 |
|  | AgaguU | 7.89 | 5.82 | 7.87 | 7.38 | 5.45 | 11.72 | 55 / 45 ± 3 |
|  | UgaguA |  |  |  |  |  | 12.03 | 42 / 58 |
|  | CgaguG | 7.7 | 5.63 |  | 7.56 |  | 10.84 | 100 / 0 |
| std dev |  | 0.221 | 0.195 | 0.014 | 0.200 | 0.057 | 0.850 |  |

The other of the two slowly exchanging conformations, or “alternative” conformations, is one of two possibilities (conformations II and III in Fig. 1) or a mixture of the two. A summary of conformation populations for each sequence is given in Table 2. Details of each sequence follows.

**Table 2.**
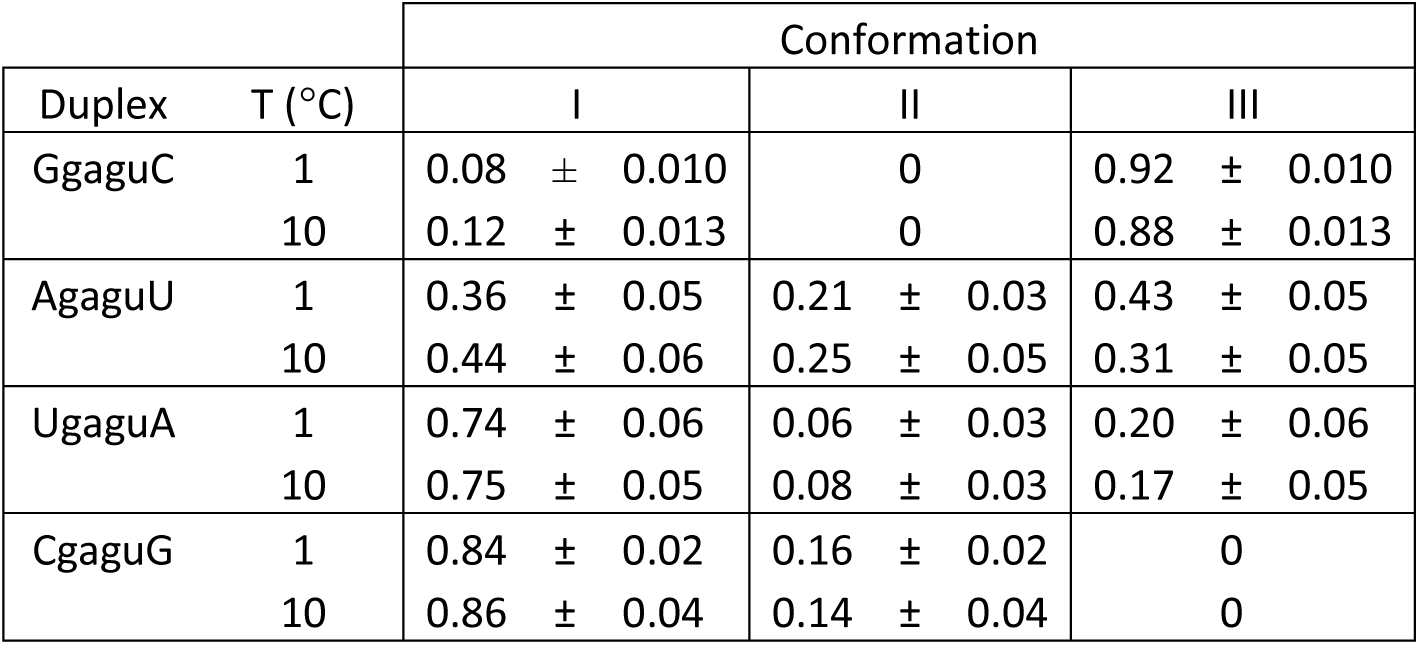
Fractions of conformations I, II and III for each duplex at 1 and 10 °C. Calculation of fractions does not include hairpin conformation or the alternative pairing in the stem of AgaguU.

| Duplex | T (°C) | Conformation |  |  |
| --- | --- | --- | --- | --- |
|  |  | I | II | III |
| GgaguC | 1 | 0.08 ± 0.010 | 0 | 0.92 ± 0.010 |
|  | 10 | 0.12 ± 0.013 | 0 | 0.88 ± 0.013 |
| AgaguU | 1 | 0.36 ± 0.05 | 0.21 ± 0.03 | 0.43 ± 0.05 |
|  | 10 | 0.44 ± 0.06 | 0.25 ± 0.05 | 0.31 ± 0.05 |
| UgaguA | 1 | 0.74 ± 0.06 | 0.06 ± 0.03 | 0.20 ± 0.06 |
|  | 10 | 0.75 ± 0.05 | 0.08 ± 0.03 | 0.17 ± 0.05 |
| CgaguG | 1 | 0.84 ± 0.02 | 0.16 ± 0.02 | 0 |
|  | 10 | 0.86 ± 0.04 | 0.14 ± 0.04 | 0 |

### CgaguG

NMR chemical shifts and NOESY data of CgaguG indicate properties similar to those observed in previous studies of this sequence . Conformation I represents 84±2% of duplexes forming the GAGU internal loop with the remainder forming an alternative conformation. A minor hairpin conformation is also observed. The alternative duplex conformation was previously identified by comparison with modified constructs and MD analysis to consist of a narrow minor groove with a single hydrogen bond in the A-G pairs (G6amino – AN1) and a bifurcated hydrogen bond in the G-U pairs (21).

Spectral properties of this alternative duplex conformation, which we refer to as conformation II, include no observable U7 imino resonance nor any cross peak to it from G4 imino as expected in a wobble GU pair. Further, the resonance for G6 imino (assigned by the exchange cross peak to G6 imino of the major conformation) is broadened by exchange with water (note lack of diagonal peak intensity, Fig. 2) and is more upfield (∼11 ppm; Table 1) relative to other tandem AG imino pairs (see GgaguC below; (17)), and lacks a NOESY cross-peak to A5*H2. However, an NOE between A5*H2 and G6 amino (G6H2) is noted (see Fig. 8 from Spasic et al. (21)), consistent with the single hydrogen bond in the AG pair of conformation II and despite the low population of conformation II in this sequence. There is no contribution from conformation I to this NOE because the A5*H2-G6H2 distance is approximately 6 Å.

### GgaguC

NMR properties of GgaguC indicate that swapping the “closing” C-G pair with a G-C pair results in G-U wobble and A-G imino pairs. We refer to this as conformation III (Fig. 1). Notably, GgaguC prefers conformation III almost exclusively over conformation I, which is populated at just 8±1% at 1 °C. G-U wobble pairs are indicated by the sharp U7H3 peak indicating this proton is stabilized by a hydrogen bond and observation of the strong U7H3- G4*H1 cross-peak typical of G-U wobble. The A-G imino pairs are indicated by the relatively sharp and downfield G6H1 signal (∼13 ppm; Table 1) and cross-peaks A5H2-G6*H1 and A5H2- G6*amino.

The duplex GgaguC is present almost entirely in a single structure, conformation III, which provides the opportunity to apply standard methods for generating restraints and applying them to model the structure. NMR restraints applied in a simulated-annealing procedure included A-G imino pairs and G-U wobble pairs each with two hydrogen-bonds resulting in the 4×4 internal loop. All other pairs are canonical base pairs. A more subtle feature includes an apparent *trans* conformation of the G6 y torsion (y ∼ 180°) instead of the A-form *g*^+^ conformation. This torsional arrangement is supported by NOEs suggesting a short G6H8-H5’’ distance and similarity to spectra of A-G tandem pairs closed with pairs other than G-U (Fig. S2). In RNA backbones a *trans*-to-*g*^+^ y transition is commonly accompanied by an alpha switch from –70° to +140°, the crankshaft conformation, also called conformation 1c (42). The G6 y torsion in GgaguC was restrained in these calculations to the *trans* conformation. Nonetheless, 22 conformers in the ensemble of 600 models violated this restraint in one strand of the duplex, suggesting substantial dynamics in the backbone for this sequence. An overlay of the lowest restraint energy structures in the resulting ensemble is shown in Fig. 3 and structure statistics are in Table S3.

**Figure 3.**
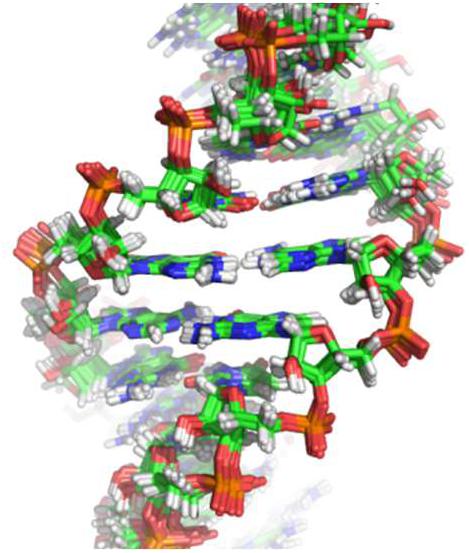
Ensemble of 10 models of GgaguC with lowest restraint energies resulting from simulating-annealing with NMR restraints.

### UgaguA and AgaguU

NMR spectra suggest that UgaguA and AgaguU have properties that are a mixture of conformation II and III when not in conformation I. Of the portions of these duplexes that are either I, II, or III, UgaguA and AgaguU are found to be approximately 74% and 36%, respectively, in conformation I. The remaining 26% and 64%, respectively, exist in alternative conformations – a mixture of II and III. The mixture is supported by the observation that chemical shifts of protons in the central A-G pairs are intermediate to the shifts for conformation II (CgaguG) and conformation III (GgaguC). These trends, shown in Fig. 4 and quantified as populations of II and III in Tables 1 and 2, indicate a preference for conformation III. Further, each of these sequences indicates a weak U7H3-G4*H1 NOE cross-peak (Fig. 2) suggestive of a G-U wobble pair with the U imino stabilized by a hydrogen bond. No evidence of A5H2-G6*H1 is noted as expected in conformation III, but G6H1 is very broad in both sequences, which would likely prevent observation of this NOE. However, the related A5H2-G6*amino NOE is clearly evident, with intensities more similar to those expected for conformation III than the weaker intensities expected for conformation II and observed for CgaguG (Fig. 2, insets). The observation of just one resonance for the alternative conformation suggests that conformations II and III are in fast exchange (microsecond range) (43,44). Apparently, this exchange is slower for UgaguA than for AgaguU as two peaks are observed for UgaguA G6H1 (Fig. S3), which is the proton with the largest chemical shift change between conformation II and III (Table 1). The rate of exchange between conformation I and the alternative conformation in AgaguU is estimated at 25-50 ms based on 2D exchange spectra acquired as a function of mixing time (Fig. S1; inset), similar to that observed previously for CgaguG (21). In addition to conformation I, conformation II/III, and hairpin, AgaguU forms a minor population that likely involves an alternative pairing in the stem that includes a GU wobble pair (Fig. 2, peaks at 11.6 and 11.9 ppm).

**Figure 4.**
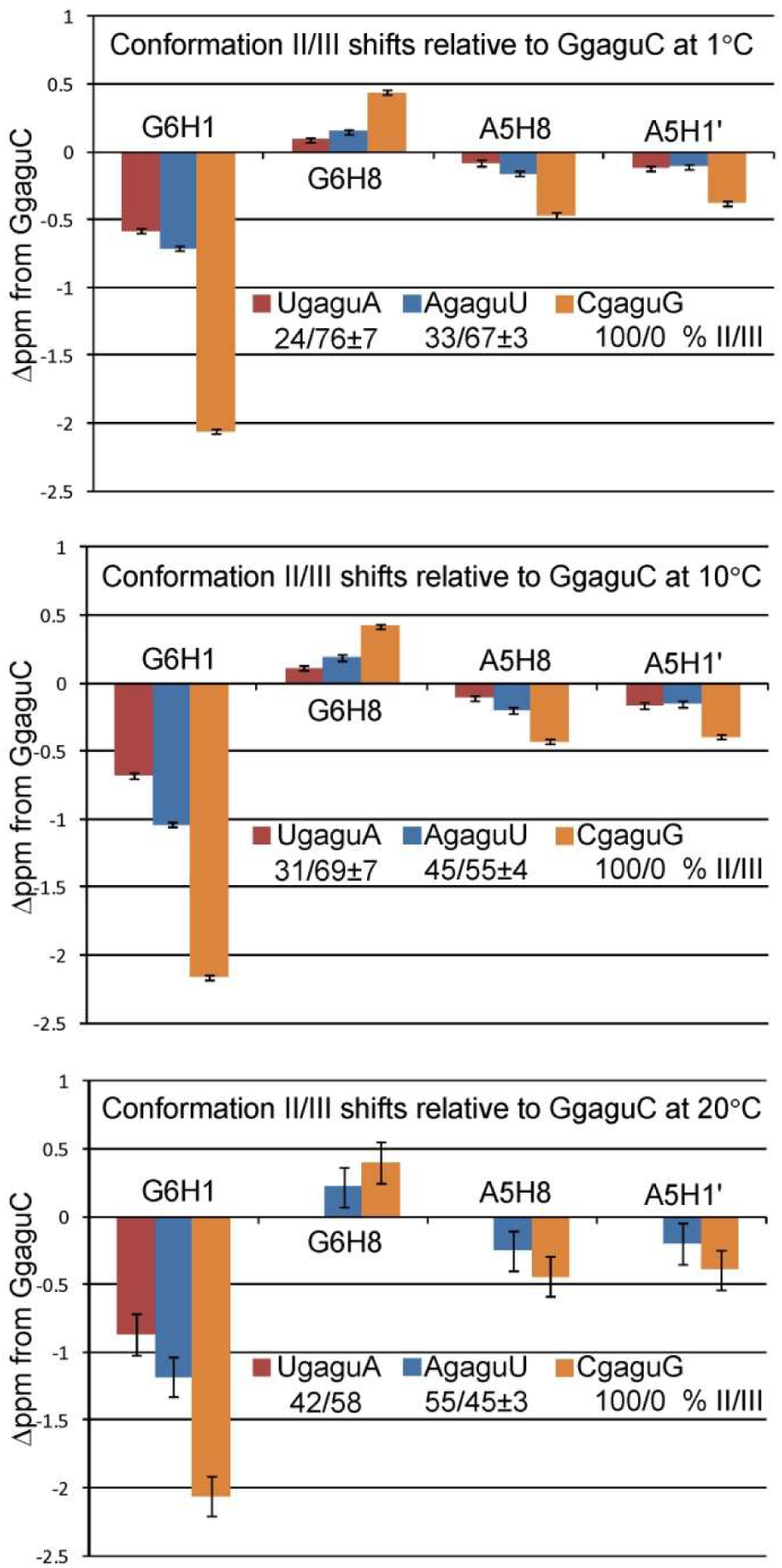
Conformation II/III chemical shift differences of assigned A5 and G6 protons in UgaguA, AgaguU and CgaguG relative to GgaguC at 1, 10, and 20 °C. A5 and G6 atoms with a difference less than 0.2 ppm between CgaguG and GgaguC are not shown. With the assumption that conformation II and III are the only alternative conformations present, percent of conformation II or III are calculated from average of A5H8, A5H1’, G6H8, and G6H1 and variance of those averages assuming that CgaguG represents 100% conformation II and GgaguC represents 100% conformation III.

### Dynamic properties and exchange between conformations II and III

Linewidths of loop protons are generally two to three-fold narrower in conformation I than in the alternative conformations (Table 3). This suggests that while conformation I is well-defined, the alternative conformations are sampling multiple conformations. In some cases the large linewidths could be due to exchange on an intermediate time-scale between conformation II and III. For instance, the ∼2 ppm chemical shift change of G6H1 between conformations II and III explains line-widths >100 Hz in the loops with A-U and U-A closing pairs if the two conformations are comparably populated with exchange time of 10-100 microseconds. The loops with G-C and C-G closing pairs are not expected to show II to III exchange broadening because in each case one conformation dominates. On the other hand, the width of G6H8 in conformation III (GgaguC) appears to be twice that for conformation II (CgaguG), suggesting a dynamic process not present in conformation II. G6H8 linewidths in the loops with A-U or U-A closing pairs are intermediate to linewidths noted for conformation II and III. Some of the G6H8 broadening in the loops with A-U or U-A adjacent pairs may be due to II to III exchange (much less than for G6H1 because G6H8 changes by just 0.44 ppm going from III to II). However, we interpret this broadening to be caused by the extra dynamic process in III, consistent with the idea that these sequences contain a rapidly exchanging mixture of II and III.

**Table 3.** Spectral widths (Hz) of aromatic and G6 imino protons in each sequence at 1 °C of conformation I and conformation II/III. Column P(3/8)H8 refers to H8 of G3, A3, A8, or G8 depending on sequence.

| 1 °C |  | G6H1 | G6H8 | A5H2 | A5H8 | G4H8 | P(3/8)H8 |
| --- | --- | --- | --- | --- | --- | --- | --- |
| Conformation I | GgaguC | 23 ± 2.6 | 10 ± 0.4 | 11 ± 0.5 | 11 ± 1.2 |  |  |
|  | AgaguU | 21 ± 1.6 | 8 ± 0.7 | 6 ± 0.7 | 8 ± 1.1 | 12 ± 0.4 | 9 ± 1.3 |
|  | UgaguA | 20 ± 0.7 | 8 ± 0.5 | 7 ± 0.5 | 8 ± 0.4 | 9 ± 1.6 | 9 ± 0.6 |
|  | CgaguG | 17 ± 0.3 | 8 ± 0.8 | 7 ± 0.3 | 8 ± 0.7 | 11 ± 0.5 | 9 ± 0.3 |
| Conformation II/III | GgaguC | 46 ± 2.2 | 44 ± 7.9 | 21 ± 2.6 | 20 ± 0.3 | 11 ± 0.3 | 10 ± 0.4 |
|  | AgaguU | >200 | 40 ± 1.6 | 19 ± 1.3 | 23 ± 1.2 | 16 ± 2.2 | 10 ± 1.0 |
|  | UgaguA | >150 | 42 ± 2.4 | 24 ± 2.7 | 22 ± 1.0 |  | 19 ± 1.7 |
|  | CgaguG | 53 ± 6.4 | 19 ± 2.1 | 25 ± 0.9 | 23 ± 0.9 | 14 ± 1.2 | 18 ± 0.6 |

### MD simulations

Each sequence and structure was simulated in four replicates for 1 μs each. A set of simulations was run at 10 °C and in 0.1 M K^+^ to match the NMR experiments and another set was run at 10 °C, but in 1 M K^+^ to test whether the conformations depended on counterion concentration. **Table 4** summarizes the simulations with respect to sequence, starting structure, K^+^ concentration, number of replicates, and simulation time for each replicate. The root mean squared deviation (RMSD) with respect to the energy-minimized structure for each replicate of each sequence is plotted in **Fig. S4**. The plots indicate that the helical regions stayed intact with average RMSD of about 1 Å for all trajectories. The RMSD values for complete structures (helices and loop) reflected a conformational variability driven by the flexibility of the loop regions. Simulations starting in conformation I were generally stable with whole-structure average RMSD not exceeding 1.8 Å for all sequences except for replicate four of GgaguC in 0.1 M K^+^, which had an average value of 4.7 Å. Simulations starting in conformation III had higher average whole-structure RMSD values in CgaguG and UgaguA. This was as high as 4.6 Å for the average of replicate 1 of UgaguA. In addition, the RMSD plot of replicate 2 of UgaguA suggested interchange between two conformations.

**Table 4.**
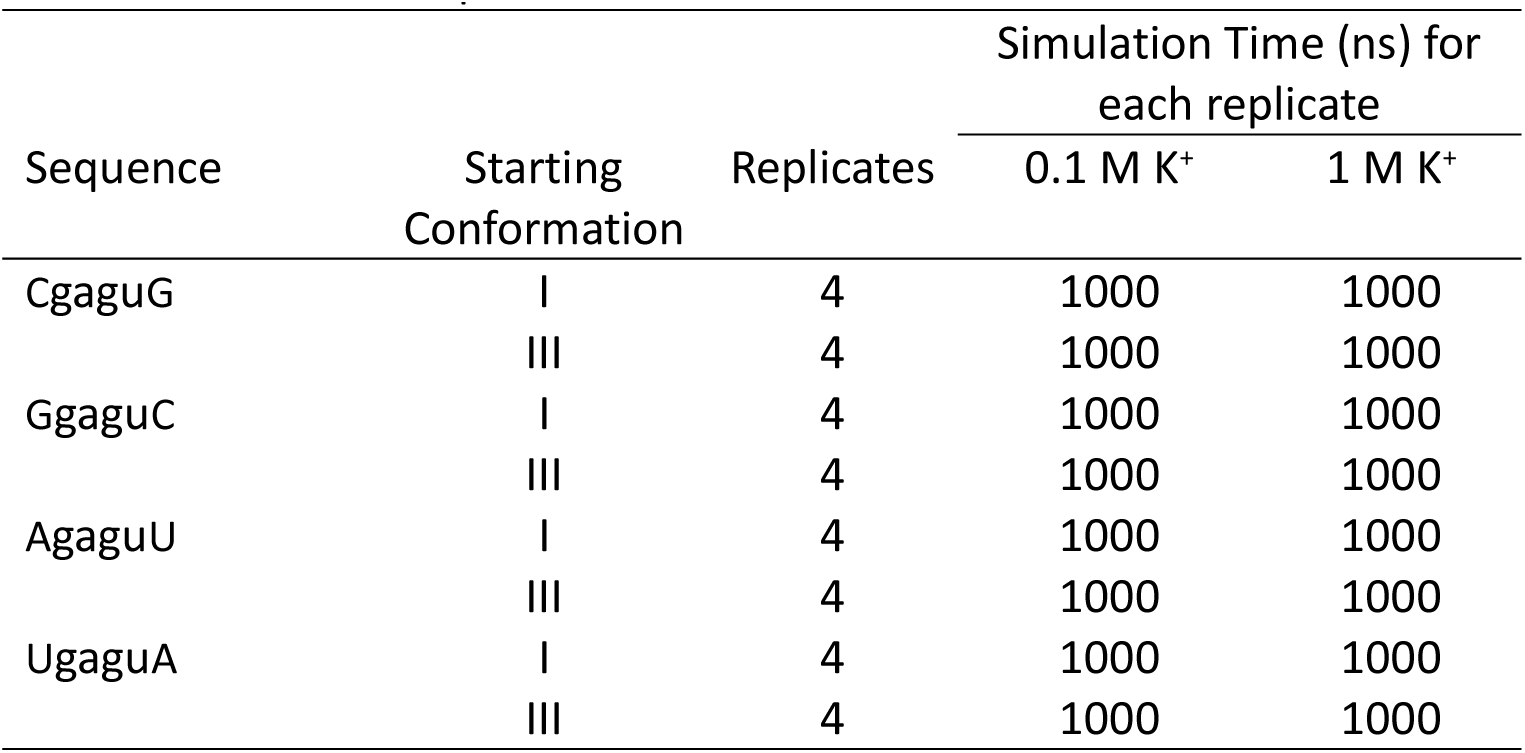
A total of 64 simulations were run (4 sequences, 2 starting conformations, 2 solvent molarities and 4 replicates). The total simulation time is 64 μs.

| Sequence | Starting Conformation | Replicates | Simulation Time (ns) for each replicate |  |
| --- | --- | --- | --- | --- |
|  |  |  | 0.1 M K<sup>+</sup> | 1 M K<sup>+</sup> |
| CgaguG | I | 4 | 1000 | 1000 |
|  | III | 4 | 1000 | 1000 |
| GgaguC | I | 4 | 1000 | 1000 |
|  | III | 4 | 1000 | 1000 |
| AgaguU | I | 4 | 1000 | 1000 |
|  | III | 4 | 1000 | 1000 |
| UgaguA | I | 4 | 1000 | 1000 |
|  | III | 4 | 1000 | 1000 |

### Clustering of conformations from simulations

Trajectory replicates of each sequence were concatenated and then clustered by conformations. **Table 5** shows the fractional composition of each cluster and the conformation of the centroid structure in the cluster for 0.1 M and 1.0 M K^+^ simulations. Across the sequences, higher K^+^ tended to increase the population of conformation III, with UgaguA being an exception to this.

**Table 5.** Clustering analysis of simulations in 0.1 M and 1.0 M K^+^. All four replicates were concatenated prior to clustering. Conformation of the centroid of each cluster is in parentheses following the cluster population. “Inter” represents a conformation that is intermediate between II and III. These results can be compared to the NMR data in Table 2.

| Clustering Statistics |  |  |  |  |  |
| --- | --- | --- | --- | --- | --- |
|  |  | Starting Conformation I |  | Starting Conformation III |  |
| Sequence | Cluster # | 0.1 M K<sup>+</sup> | 1 M K<sup>+</sup> | 0.1 M K<sup>+</sup> | 1 M K<sup>+</sup> |
| CgaguG | a | 1.00 (I) | 1.00 (I) | 0.92 (II) | 0.75 (II) |
|  | b | - | - | 0.08 (III) | 0.25 (III) |
| AgaguU | a | 1.00 (I) | 1.00 (I) | 1.00 (II) | 0.90 (II) |
|  | b | - | - | - | 0.10 (III) |
| UgaguA | a | 1.00 (I) | 1.00 (I) | 0.39 (II) | 0.30 (III) |
|  | b | - | - | 0.36 (III) | 0.51 (II) |
|  | c | - | - | 0.25 (Inter) | 0.19 (Inter) |
| GgaguC | a | 0.78 (I) | 1.00 (I) | 1.00 (II) | 0.93 (II) |
|  | b | 0.22 (Inter) | - | - | 0.07 (III) |

### Simulations starting in conformation III

As seen in past simulations of CgaguG (21), for sequences starting in conformation III in the 0.1 M K^+^ simulations, the dominant population cluster was conformation II, characterized by G-U pairs with bifurcated hydrogen bonds and tandem A-G pairs with single hydrogen-bonds (Fig. 1). Conformation III structure populations for CgaguG and UgaguA were 8 and 36%, respectively (Table 5). Conformation III had no representation for simulations of AgaguU and GgaguC; instead, conformation II was populated (Table 5).

Simulations of UgaguA populated intermediate state conformations, as shown in Fig. S5, with G4-U7* (and G4*-U7) and A5-G6* (and A5*-G6) pairs having III and II-like orientation, respectively. Simulations starting at conformation III estimate that CgaguG populated conformation II at 92%, in general agreement with NMR (Tables 2 and 5). Simulations for UgaguA estimate conformation II and III are approximately equally populated, while NMR indicates approximately a 2:1 (10 °C) or 3:1 (1 °C) preference for conformation III. MD simulations of AgaguU and GgaguC, which estimate almost no occurrence of conformation III, have the greatest deviation from the NMR observations. NMR indicates AgaguU has a 1:1 (10 °C) or 2:1 (1 °C) preference for conformation III:II and that GgaguC exists only in conformation III (Table 1). G6 and G6* alpha, beta, gamma and delta dihedral angles of GgaguC closely match those of A-form (42) (Table S4), which is inconsistent with the crankshaft conformation observed in 2D NOE of the sequence (NMR Results above, section GgaguC).

### Simulations starting in conformation I

Simulations starting in conformation I did not transition into II or III. Structures clustered into a single cluster for all sequences except GgaguC (Fig. 5), for which 22% of structures were in a cluster (Fig. S5) characterized by a C3’endo sugar pucker for G4 and G4*. This cluster, however, existed only in replicate 4 (Fig. S6).

**Figure 5.**
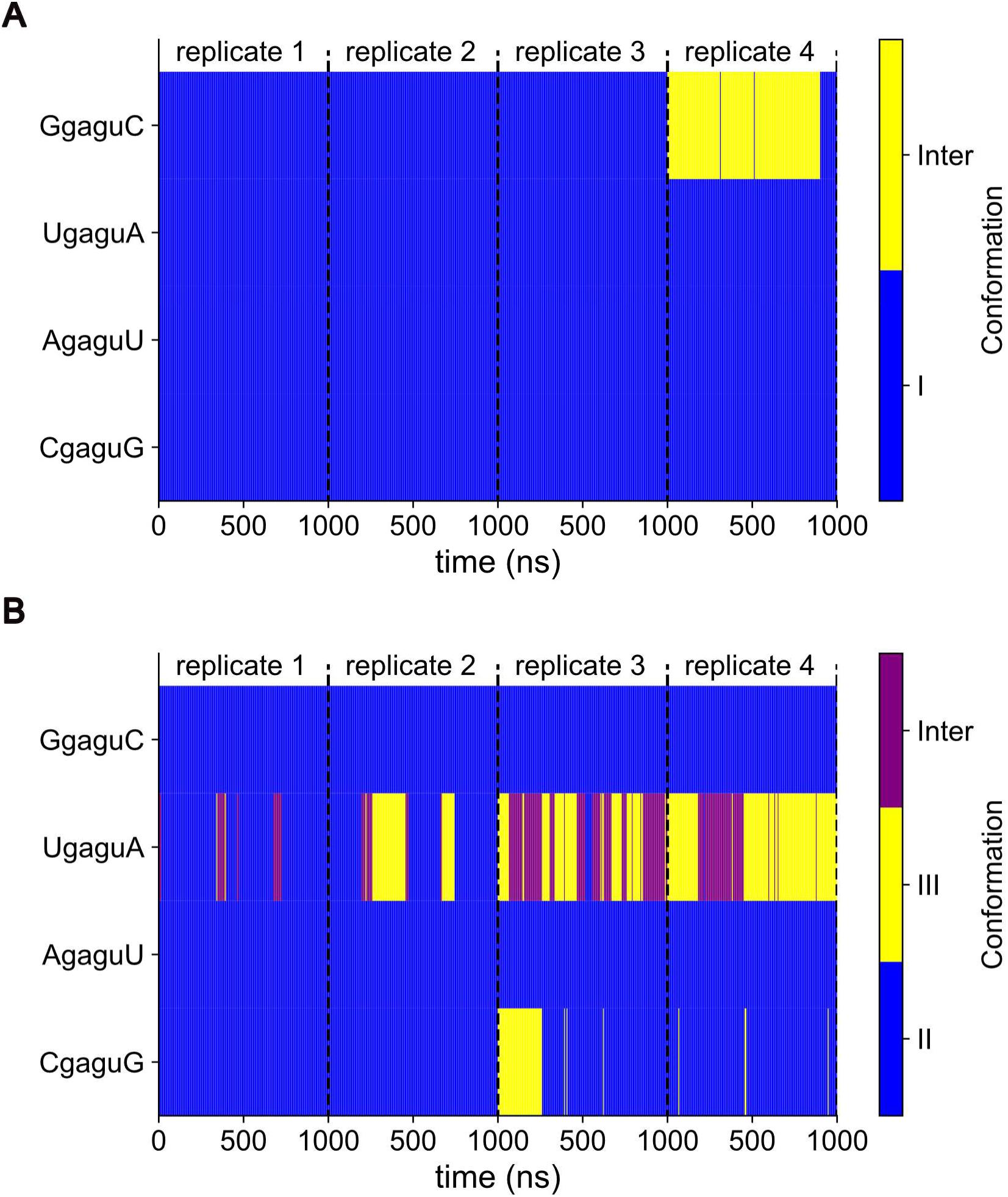
Time series plot of cluster index for structures generated in MD simulation (0.1 M K^+^). Simulations starting in conformation I (panel A) showed clear signs of incomplete convergence marked by the presence of intermediate structures in replicate four of GgaguC. Similarly, in simulations starting in conformation III (panel B), there was a fairly large population of structures in conformation III observed in replicate 3 of CgaguG not present to much extent in the other 3 replicates of the sequence. In addition, UgaguA suggested interchange between three conformations.

Fig. 6 shows the χ dihedral angles as a function of time for residues in the G4/G6* base pairs of simulations starting in conformation I. G4 and G4* consistently populated the *syn* starting conformation (χ ∼ 0 - 90°). In AgaguU and CgaguG, G6 and G6* presented as both *syn* and *anti* (χ ∼ 90 – 270°). In GgaguC and UgaguA, G6 and G6* χ remained largely in the *syn* conformation, although in 1 M K^+^ GgaguC and UgaguA make transitions to G6/G6* *anti* (Fig. S7). The reduced population of G6/G6* *syn* conformation relative to G4/G4* is consistent, although not in quantitative agreement, with NMR evidence based on the relative intensities of H8-H1’ NOE cross-peaks. The G6/G4 H8-H1’ 1/r^6^ ratio was calculated for the concatenated trajectory of each system and compared to the NMR results (Table S2). This ratio was 0.54 and 0.59 for CgaguG and AgaguU, respectively, for the 0.1 M K^+^ simulation. UgaguA and GgaguC had a ratio of 1.03, consistent with Fig. 6 where there were no substantial transitions between *syn* and *anti* conformations for these systems. In the 1.0 M K^+^ simulations, however, all four systems had a G6/G4 H8-H1’ 1/r^6^ ratio less than 1 with CgaguG, AgaguU, UgaguA and GgaguC having 0.31, 0.51, 0.80 and 0.94, respectively, similar to NMR observations. The instability of the χ angle distribution of G6 and G6*, in contrast to that of G4 and G4*, might be a result of the former’s contiguity to U7 and U7*, which is flipped out of the structure and in the solvent.

**Figure 6.**
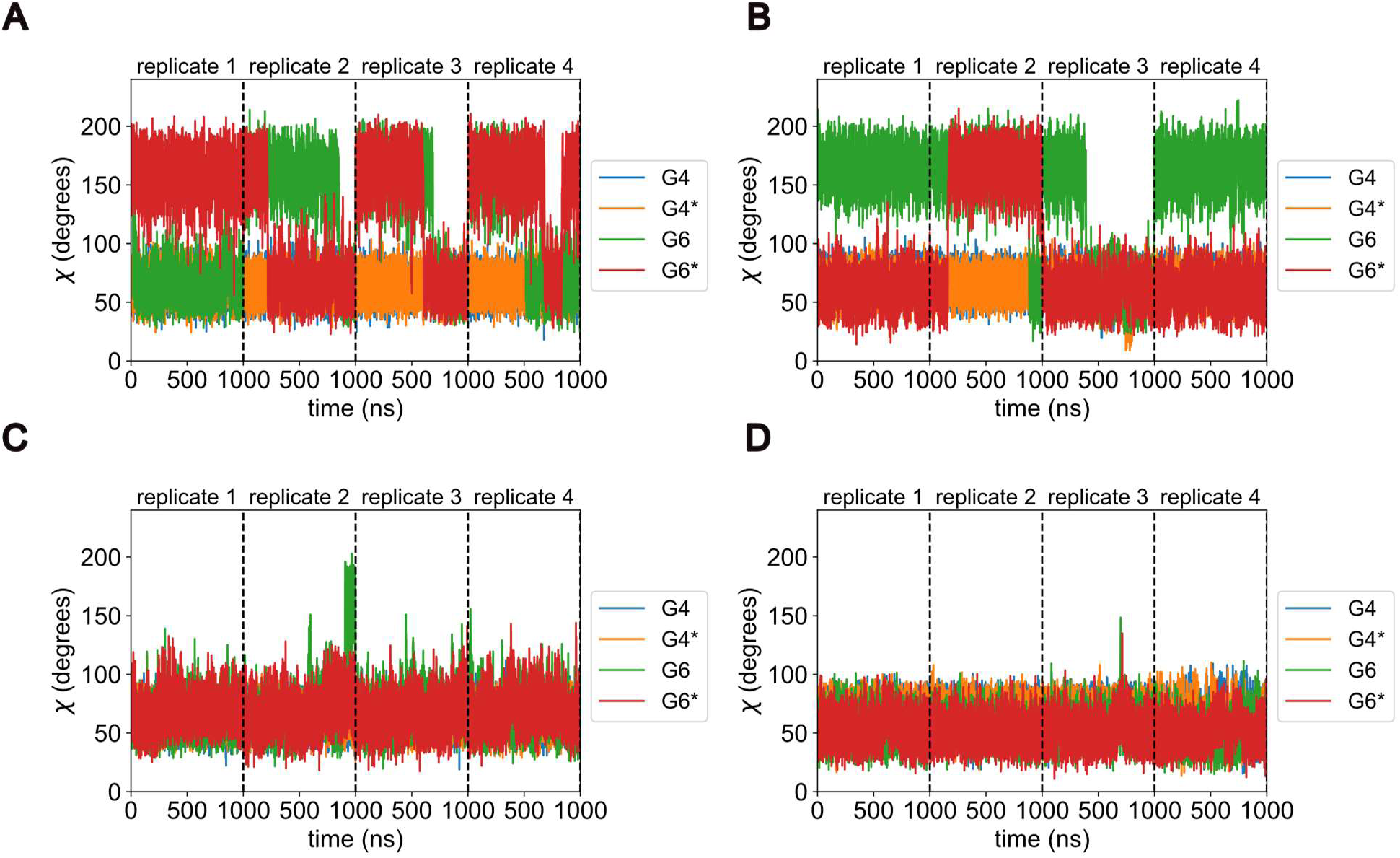
χ dihedral angles as a function of simulation time for G4, G4*, G6 and G6* for simulations in 0.1 M K^+^ starting in conformation I. Labels rep1, rep2, rep3 and rep4 refer to replicate trajectories 1, 2, 3 and 4, respectively. The vertical broken lines represent the end of a replicate and/or the start of the next.

### 1M K^+^ Simulations

Simulations in 1 M K^+^, much like those in 0.1 M K^+^, had most of the structures in conformation II for all sequences starting in conformation III (Table 5). In addition, all sequences, except GgaguC, starting in conformation I were clustered into a single group, similar to simulations in 0.1 M K^+^, with no transitions into II/III conformations. Unlike the 0.1 M K^+^, however, simulations of AgaguU and GgaguC starting in conformation III in 1 M K^+^ had low populations (10% and 7%, respectively) of structures in a cluster consistent with conformation III, while all the structures of these sequences in lower K^+^ simulations were in a single cluster consistent with conformation II (Table 5). Also, CgaguG showed a modest increase of conformation III in 1 M K^+^ (25%, up from 8% in 0.1 M K^+^). The intermediate state observed for UgaguA structures in 0.1 M K^+^ was similarly present in 1 M K^+^, with conformation III-like G-U pairs and II-like A-G pairs. Finally, as discussed above, the reduced population of G6/G6* syn conformation in AgaguU and CgaguG in the 0.1 M K^+^ simulations was reproduced in all four duplexes in 1 M K^+^.

### Convergence

Given that the conformational populations of conformations II and III did not agree well with the NMR data, we tested convergence by clustering both the first half and full-length trajectories of each simulation. For acceptable convergence given our starting conformations and four replicates of 1 μs simulations, we expect that cluster populations would not deviate considerably between 0.5 μs and 1 μs.

Fig. 7 (for 0.1 M K^+^) and Fig. S8 (for 1 M K^+^) show the population abundance values of clusters with centroids in conformations II and III for half and full-length simulations averaged over the four replicates of each sequence. For conformation II/III structures, extending the simulation from 500 ns to 1 µs did not substantially perturb cluster population statistics, thus there is no evidence of a lack of convergence for these simulations. For both simulation times, structures in conformation II constitute the dominant population, even though the simulations started in conformation III.

**Figure 7.**
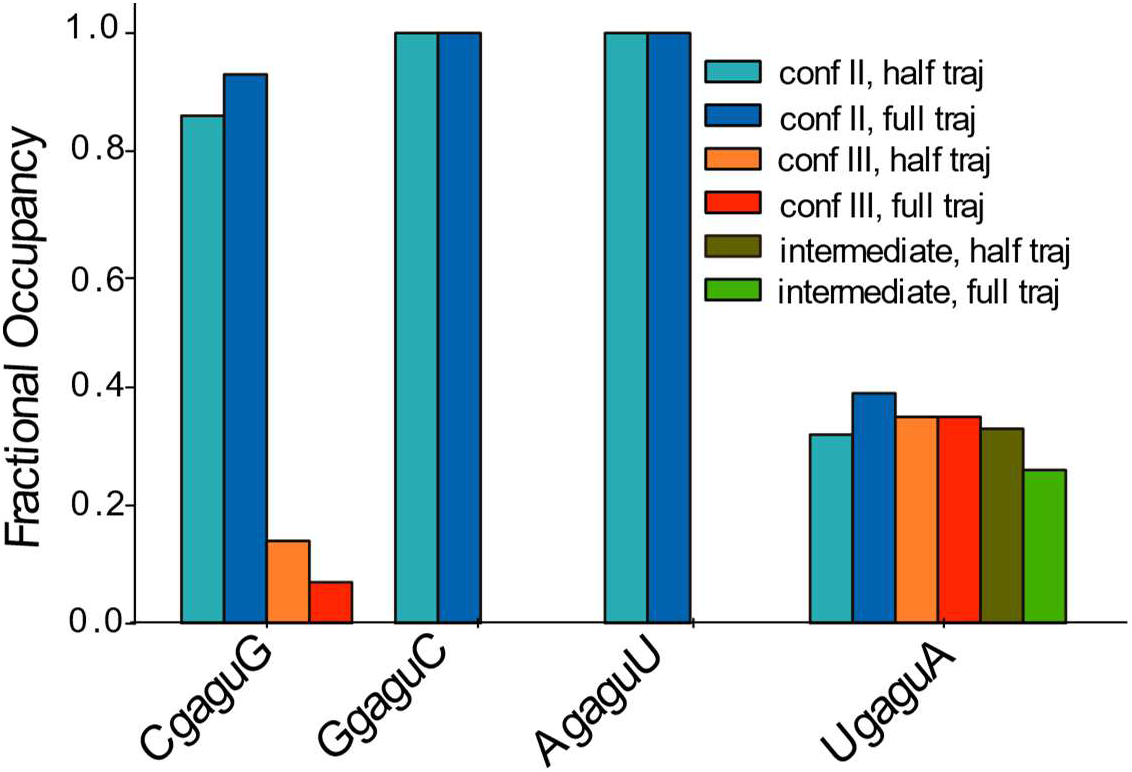
The average fractional population of clusters of trajectory structures generated in simulations starting in conformation III over four replicates in the first 500 ns and the entire 1 µs of simulation. Results show that extending the simulations from 500 ns to 1 µs does not substantially alter cluster statistics. Duplex UgaguA has a third cluster of structures with intermediate conformation.

In general, the population of structures in conformation II marginally increased in the second half of the full trajectories. This was because trajectories needed to transition from conformation III to II, which occurred in the first 500 ns for most replicates of the sequences. There were three exceptions: UgaguA in 1 M K^+^, which had a substantial population of intermediates, and AgaguU and GgaguC in 0.1 M K^+^, which occupied a single cluster (conformation II) across the full 1 μs simulations.

We also followed the conformation as a function of time across the trajectories (0.1 M K^+^ Fig. 5 and 1 M K^+^ Fig. S9). Simulations starting in conformation I did present evidence that the simulations were not fully converged. Replicate 4 of GgaguC in 0.1 M K^+^ and replicate 3 in 1 M K^+^ contained conformations that were not observed in all four replicates of the simulations (C3’endo sugar pucker for G4 and G4*), despite being the dominant structural conformation in the replicates in which they were present. **Fig. 7 and Fig. S7** show that the χ backbone fluctuations of G6 and G6* were generally not converged in simulations in 0.1 and 1 M K^+^, respectively. As noted above, this is possibly due to their proximity to the high-entropy U7 and U7* nucleotides.

## DISCUSSION

RNA systems that populate multiple conformations provide challenging tests of molecular dynamics force fields. The 4×4 internal loop 5’GAGU/3’UGAG populates three conformations, although in previous work only two unusual conformations (I and II) were experimentally detected in the sequence CgaguG (Fig. 1) (15,21). The expected conformation (III with A-G imino pairs) based on prior studies for tandem A-G pairs with canonical closing pairs was not observed (16–20), except that replacement of G6 with inosine or G-locked nucleic acid (LNA) resulted in conformation III (21). The NMR experiments presented here show that if the C-G pairs that close the 4×4 loop are changed to G-C, A-U, or U-A, populations of A-G imino / G-U wobble pairs are detected (conformation III). The sequences all differ in the relative populations of I, II, and III.

In NMR, a separate resonance is observed for conformation I in all sequences. Exchange between I and the alternative conformations, II and/or III, is indicated by exchange cross-peaks in the NOESY spectra; these can only appear if exchange is occurring on a timescale on the order of the mixing time and faster than longitudinal relaxation (∼1 sec) (45). Thus, exchange between I and the alternative conformations is likely on the order of 5-200 ms, slow enough for observation of separate peaks and fast enough to occur during the mixing time. Direct measurement of exchange cross-peak evolution as a function of mixing time in AgaguU and CgaguG demonstrated that, in these two sequences, exchange between I and the alternative conformation occurs on the order of 25-50 ms (Fig. S1; (21)). Apparently the choice of closing pair does not substantially alter the slow exchange property of conformation I with II/III. The interchange between conformations II and III happens on the order of milliseconds or faster, for the most part, as only one NMR peak is observed for these conformations (43,44). G6H1 of UgaguA is the exception, suggesting slower II-III exchange for this sequence.

### Comparison of Simulations to NMR

Molecular dynamics simulations using the latest Amber force field (22–24) starting at either conformation I or III reflect many of the properties observed experimentally. Simulations of all sequences starting as conformation I primarily remain as I, with few transitions to other conformations and no transitions to either II or III (Table 5). Because the simulations are only a microsecond in duration, this agrees with the millisecond time scale observed experimentally for transitions from I to the alternative conformations.

Additionally, a conformational change in the backbone in most simulations starting in conformation I results in G6 χ changing from *syn* to an *anti* orientation, consistent with the G6H8-H1’ NOESY cross-peak being weaker than the equivalent cross-peak of G4 in at least three duplexes. The χ of G4 never leaves a *syn* orientation in MD simulations. The instability of the G6 and G6* χ angle distribution, in contrast to stable *syn* chi in G4 and G4*, may be a result of the former’s contiguity to U7 and U7* which are flipped out of the structure and in the solvent.

Simulations starting in conformation III do not show transitions to I, but multiple transitions to II-like clusters and back are observed (Fig. 5). Notably, few clusters other than II or III are visited.

MD simulations cannot be used to predict the relative stability of I vs II/III because there are no transitions between these states. Sufficient transitions, however, between conformations II and III occur to reach conclusions about relative stability of II vs III, which can be compared to NMR experiments (Fig.s 5 and S9). While NMR and MD results agree that conformation II is dominant in CgaguG, they do not agree well in the other three sequences, especially in GgaguC and AgaguU, where NMR indicates dominance of conformation III while MD suggests almost none of that conformation. MD predicts substantial populations of conformation III in UgaguA although not to the extent suggested by NMR.

### MD Demonstrates Intermediate Structures

The presence of multiple II to III transitions allows the possibility of examining intermediate structures. The A-G pair of the UgaguA intermediate in Fig. S5 exhibits the Watson-Crick-Franklin face of G rotating slightly away from the Watson-Crick-Franklin face of A, resulting in replacement of the GH1-AN1 hydrogen bond with an apparently bifurcated hydrogen bond between the G amino protons and AN1. GH1 remains somewhat protected from solvent unlike conformation II, but overall this shift is in the direction of conformation II. Meanwhile, the G-U wobble pairs of conformation III remain largely intact in this intermediate, including the interaction of UH3 with GO6 which should protect UH3 from exchange with solvent. If this intermediate represents a transition state, then apparently the transition from III to II begins with a shift at the A-G imino pair, followed later by a fold which narrows the groove and breaks the G-U wobble pair.

### Importance of improved force fields

Here we studied the standard Amber force field (22,23). There are alternative force fields, but they have been found to have limitations (46–52). We hypothesized that Amber would stabilize the native conformations for GAGU because our prior simulations stabilized native conformations and helped solve the solution structure of the minor form (II) (21). This study, however, reveals the limitations of the force field. We observed an over-stabilization of conformation II for closing pairs that NMR demonstrated a preference for conformation III. This system makes an excellent additional test of force fields because the closing pairs affect the conformational choice of the loop and we can sufficiently sample conformational choice between states II and III. We expect that a good force field would match experimentally-determined populations of II and III.

### The asymmetry of GU and UG pairs and influence of conformational choice

It has been previously noted that G-U pairs entering a helix are more frequently observed with the G at the 5’ end of the helix and the U at the 3’ end of the other strand (53). This apparently is the result of undertwisting between the G-U pair and the following pair (54). The undertwisting increases the stacking of the G-U with the following pair (55).

The GAGU sequence studied here results in a U-G pair entering the adjacent helices (in conformations II and III), with the U at the 5’ end of the helix and the G at the 3’ end. This orientation of the G-U pair (counter to the preferred orientation) results in overtwisting to the following pair and less stacking. This helical twist induced by U-G wobble pairing, at least in part, explains why CgaguG and UgaguA prefer conformation I, but loops composed of U<u>AG</u>G (the preferred wobble orientation for entering a helix), A<u>AG</u>U, and G<u>AG</u>C form 2×2 internal loops with maximal (canonical and non-canonical) pairing (like that in conformation III) (15–17).

GgaguC and AgaguU have an opposite orientation (with respect to purine and pyrimidine) of the pair following the U-G to CgaguG and UgaguA. This orientation of the next pair alters the conformational preference from I to II/III (Table 2), increasing the pairing. Evidently the stacking on a following purine-pyrimidine pair provides energetically favorable stacking in spite of the overtwisting. These features should be considered in modeling of 3D structures and designing of 3D folds (56).

## AUTHOR CONTRIBUTIONS

OA designed computational research, performed computational research, analyzed data and wrote the manuscript. SDK designed NMR research, performed NMR research, analyzed data, and wrote the manuscript. JS provided instruction for computational research and edited the manuscript. DHM designed research and wrote the manuscript.

## ACKNOWLEDGEMENTS

This work was supported by NIH grant R35GM145283 to D.H.M. The University of Rochester Center for Integrated Research Computing provided computational resources.

## REFERENCES

1. Serganov, A., and E. Nudler. 2013. A decade of riboswitches. Cell. 152(1-2):17–24, doi: 10.1016/j.cell.2012.12.024, https://www.ncbi.nlm.nih.gov/pubmed/23332744.

2. Doudna, J. A., and T. R. Cech. 2002. The chemical repertoire of natural ribozymes. Nature. 418(6894):222–228, doi: 10.1038/418222a, https://www.ncbi.nlm.nih.gov/pubmed/12110898.

3. McManus, M. T., and P. A. Sharp. 2002. Gene silencing in mammals by small interfering RNAs. Nat Rev Genet. 3(10):737–747, doi: 10.1038/nrg908, https://www.ncbi.nlm.nih.gov/pubmed/12360232.

4. Marraffini, L. A. 2015. CRISPR-Cas immunity in prokaryotes. Nature. 526(7571):55–61, doi: 10.1038/nature15386, https://www.ncbi.nlm.nih.gov/pubmed/26432244.

5. Huang, Z. H., Y. P. Du, J. T. Wen, B. F. Lu, and Y. Zhao. 2022. snoRNAs: functions and mechanisms in biological processes, and roles in tumor pathophysiology. Cell Death Discov. 8(1):259, doi: 10.1038/s41420-022-01056-8, https://www.ncbi.nlm.nih.gov/pubmed/35552378.

6. Chauvier, A., J. Cabello-Villegas, and N. G. Walters. 2019. Probing RNA structure and interaction dynamics at the single molecule level Methods. 162(163):3–11, doi: 10.1016/j.ymeth.2019.04.002.

7. Korostelev, A., D. N. Ermolenko, and H. F. Noller. 2008. Structural dynamics of the ribosome. Curr Opin Chem Biol. 12(6):674–683, doi: 10.1016/j.cbpa.2008.08.037, https://www.ncbi.nlm.nih.gov/pubmed/18848900.

8. Zhang, H., and S. C. Keane. 2019. Advances that facilitate the study of large RNA structure and dynamics by nuclear magnetic resonance spectroscopy. Wiley Interdiscip Rev RNA. 10(5):e1541, doi: 10.1002/wrna.1541, https://www.ncbi.nlm.nih.gov/pubmed/31025514.

9. Rangadurai, A., E. S. Szymaski, I. J. Kimsey, H. Shi, and H. M. Al-Hashimi. 2019. Characterizing micro-to-millisecond chemical exchange in nucleic acids using off-resonance R(1rho) relaxation dispersion. Prog Nucl Magn Reson Spectrosc. 112–113:55-102, doi: 10.1016/j.pnmrs.2019.05.002, https://www.ncbi.nlm.nih.gov/pubmed/31481159.

10. Zhang, K., and A. T. Frank. 2021. Probabilistic Modeling of RNA Ensembles Using NMR Chemical Shifts. J Phys Chem B. 125(35):9970–9978, doi: 10.1021/acs.jpcb.1c05651, https://www.ncbi.nlm.nih.gov/pubmed/34449236.

11. Bonilla, S. L., and J. S. Kieft. 2022. The promise of cryo-EM to explore RNA structural dynamics. J Mol Biol. 434(18):167802, doi: 10.1016/j.jmb.2022.167802, https://www.ncbi.nlm.nih.gov/pubmed/36049551.

12. Korostelev, A. A. 2022. The Structural Dynamics of Translation. Annu Rev Biochem. 91:245–267, doi: 10.1146/annurev-biochem-071921-122857, https://www.ncbi.nlm.nih.gov/pubmed/35287473.

13. Smith, L. G., J. Zhao, D. H. Mathews, and D. H. Turner. 2017. Physics-based all-atom modeling of RNA energetics and structure. WIREs RNA. 8, e1422(5), doi: 10.1002/wrna.1422, http://www.ncbi.nlm.nih.gov/pubmed/28815951.

14. Sponer, J., G. Bussi, M. Krepl, P. Banas, S. Bottaro, R. A. Cunha, A. Gil-Ley, G. Pinamonti, S. Poblete, P. Jurecka, N. G. Walter, and M. Otyepka. 2018. RNA Structural Dynamics As Captured by Molecular Simulations: A Comprehensive Overview. Chem Rev. 118(8):4177–4338, doi: 10.1021/acs.chemrev.7b00427, http://www.ncbi.nlm.nih.gov/pubmed/29297679.

15. Kennedy, S. D., R. Kierzek, and D. H. Turner. 2012. Novel conformation of an RNA structural switch. Biochemistry. 51(46):9257–9259, doi: 10.1021/bi301372t, http://www.ncbi.nlm.nih.gov/pubmed/23134175.

16. Wu, M., J. SantaLucia, Jr., and D. H. Turner. 1997. Solution structure of (rGGCAGGCC)2 by two-dimensional NMR and the iterative relaxation matrix approach. Biochemistry. 36(15):4449–4460, doi: 10.1021/bi9625915, http://www.ncbi.nlm.nih.gov/pubmed/9109652.

17. Hammond, N. B., B. S. Tolbert, R. Kierzek, D. H. Turner, and S. D. Kennedy. 2010. RNA internal loops with tandem AG pairs: the structure of the 5’GAGU/3’UGAG loop can be dramatically different from others, including 5’AAGU/3’UGAA. Biochemistry. 49(27):5817–5827, doi: 10.1021/bi100332r, http://www.ncbi.nlm.nih.gov/pubmed/20481618.

18. Zhao, C., K. R. Rajashankar, M. Marcia, and A. M. Pyle. 2015. Crystal structure of group II intron domain 1 reveals a template for RNA assembly. Nat Chem Biol.(11):967-972, doi: 10.1038/nchembio.1949.

19. Simon, B., P. Masiewicz, A. Ephrussi, and T. Carlomagno. 2015. The structure of the SOLE element of oskar mRNA. RNA. 21(8):1444–1453, doi: 10.1261/rna.049601.115, https://www.ncbi.nlm.nih.gov/pubmed/26089324.

20. Marcia, M., and A. M. Pyle. 2012. Visualizing group II intron catalysis through the stages of splicing. Cell. 151(3):497–507, doi: 10.1016/j.cell.2012.09.033, https://www.ncbi.nlm.nih.gov/pubmed/23101623.

21. Spasic, A., S. D. Kennedy, L. Needham, M. Manoharan, R. Kierzek, D. H. Turner, and D. H. Mathews. 2018. Molecular dynamics correctly models the unusual major conformation of the GAGU RNA internal loop and with NMR reveals an unusual minor conformation. RNA. 24(5):656–672, doi: 10.1261/rna.064527.117, http://www.ncbi.nlm.nih.gov/pubmed/29434035.

22. Perez, A., I. Marchan, D. Svozil, J. Sponer, T. E. Cheatham, 3rd, C. A. Laughton, and M. Orozco. 2007. Refinement of the AMBER force field for nucleic acids: improving the description of alpha/gamma conformers. Biophys J. 92(11):3817-3829, doi: 10.1529/biophysj.106.097782, https://www.ncbi.nlm.nih.gov/pubmed/17351000.

23. Zgarbova, M., M. Otyepka, J. Sponer, A. Mladek, P. Banas, T. E. Cheatham, 3rd, and P. Jurecka. 2011. Refinement of the Cornell et al. Nucleic Acids Force Field Based on Reference Quantum Chemical Calculations of Glycosidic Torsion Profiles. J Chem Theory Comput. 7(9):2886-2902, doi: 10.1021/ct200162x, https://www.ncbi.nlm.nih.gov/pubmed/21921995.

24. Cornell, W. D., P. Cieplak, C. I. Bayly, I. R. Gould, K. M. Merz, D. M. Ferguson, D. C. Spellmeyer, T. Fox, J. W. Caldwell, and P. A. Kollman. 1995. A second generation force field for the simulation of proteins, nucleic acids, and organic molecules. . J Am Chem Soc. 117:5179–5197, doi: 10.1021/ja00124a002.

25. Piotto, M., V. Saudek, and V. Sklenar. 1992. Gradient-tailored excitation for single-quantum NMR spectroscopy of aqueous solutions. J Biomol NMR. 2(6):661–665, doi: 10.1007/BF02192855, http://www.ncbi.nlm.nih.gov/pubmed/1490109.

26. Grzesiek, S., and A. Bax. 1993. The importance of not saturating water in protein NMR. Application to sensitivity enhancement and NOE measurements. Journal of American Chemical Society. 115:2, doi: 10.1021/ja00079a052.

27. Delaglio, F., S. Grzesiek, G. W. Vuister, G. Zhu, J. Pfeifer, and A. Bax. 1995. Nmrpipe - a multidimensional spectral processing system based on Unix pipes. J Biomol Nmr. 6(3):277–293, doi: 10.1007/BF00197809, <GO to ISI>://A1995TH72500006.

28. Lee, W., M. Tonelli, and J. L. Markley. 2015. NMRFAM-SPARKY: enhanced software for biomolecular NMR spectroscopy. Bioinformatics. 31(8):1325–1327, doi: 10.1093/bioinformatics/btu830, http://www.ncbi.nlm.nih.gov/pubmed/25505092.

29. Case, D. A., R. M. Betz, D. S. Cerutti, I. Cheatham, T.E., T. A. Darden, R. E. Duke, T. J. Giese, H. Gohlke, A. W. Goetz, N. Homeyer, S. Izadi, P. Janowski, J. Kaus, A. Kovalenko, T. S. Lee, S. LeGrand, P. Li, C. Lin, T. Luchko, R. Luo, B. Madej, D. Mermelstein, K. M. Merz, G. Monard, H. Nguyen, H. T. Nguyen, I. Omelyan, A. Onufriev, D. R. Roe, A. Roitberg, C. Sagui, C. L. Simmerling, W. M. Botello-Smith, J. Swails, R. C. Walker, J. Wang, R. M. Wolf, X. Wu, L. Xiao, and P. A. Kollman. 2016. AMBER 2016, University of California, San Francisco.

30. Still, W. C., A. Tempczyk, R. C. Hawley, and T. Hendrickson. 1990. Semianalytical treatment of solvation for molecular mechanics and dynamics. Journal of the American Chemical Society. 112:3, doi: 10.1021/JA00172A038.

31. Pettersen, E. F., T. D. Goddard, C. C. Huang, G. S. Couch, D. M. Greenblatt, E. C. Meng, and T. E. Ferrin. 2004. UCSF Chimera--a visualization system for exploratory research and analysis. J Comput Chem. 25(13):1605–1612, doi: 10.1002/jcc.20084, https://www.ncbi.nlm.nih.gov/pubmed/15264254.

32. Li, P., L. F. Song, and K. M. Merz, Jr. 2015. Systematic Parameterization of Monovalent Ions Employing the Nonbonded Model. J Chem Theory Comput. 11(4):1645–1657, doi: 10.1021/ct500918t, https://www.ncbi.nlm.nih.gov/pubmed/26574374.

33. Izadi, S., R. Anandakrishnan, and A. V. Onufriev. 2014. Building Water Models: A Different Approach. J Phys Chem Lett. 5(21):3863–3871, doi: 10.1021/jz501780a, https://www.ncbi.nlm.nih.gov/pubmed/25400877.

34. Machado, M. R., and S. Pantano. 2020. Split the Charge Difference in Two! A Rule of Thumb for Adding Proper Amounts of Ions in MD Simulations. J Chem Theory Comput. 16(3):1367–1372, doi: 10.1021/acs.jctc.9b00953, https://www.ncbi.nlm.nih.gov/pubmed/31999456.

35. Salomon-Ferrer, R., A. W. Gotz, D. Poole, S. Le Grand, and R. C. Walker. 2013. Routine Microsecond Molecular Dynamics Simulations with AMBER on GPUs. 2. Explicit Solvent Particle Mesh Ewald. J Chem Theory Comput. 9(9):3878-3888, doi: 10.1021/ct400314y, https://www.ncbi.nlm.nih.gov/pubmed/26592383.

36. Loncharich, R. J., B. R. Brooks, and R. W. Pastor. 1992. Langevin dynamics of peptides: the frictional dependence of isomerization rates of N-acetylalanyl-N’-methylamide. Biopolymers. 32(5):523–535, doi: 10.1002/bip.360320508, https://www.ncbi.nlm.nih.gov/pubmed/1515543.

37. Gomez, Y. K., A. M. Natale, J. Lincoff, C. W. Wolgemuth, J. M. Rosenberg, and M. Grabe. 2022. Taking the Monte-Carlo gamble: How not to buckle under the pressure! J Comput Chem. 43(6):431–434, doi: 10.1002/jcc.26798, https://www.ncbi.nlm.nih.gov/pubmed/34921560.

38. Ryckaert, J. P., G. Ciccotti, and H. J. C. Berendsen. 1977. Numerical integration of the cartesian equations of motion of a system with constraints: molecular dynamics of n-alkanes. J Comput Phys. 23:327–341, doi: 10.1016/0021-9991(77)90098-5.

39. Roe, D. R., and T. E. Cheatham, 3rd. 2013. PTRAJ and CPPTRAJ: Software for Processing and Analysis of Molecular Dynamics Trajectory Data. J Chem Theory Comput. 9(7):3084-3095, doi: 10.1021/ct400341p, https://www.ncbi.nlm.nih.gov/pubmed/26583988.

40. Wolf, A., and K. N. Kirschner. 2013. Principal component and clustering analysis on molecular dynamics data of the ribosomal L11.23S subdomain. J Mol Model. 19(2):539–549, doi: 10.1007/s00894-012-1563-4, https://www.ncbi.nlm.nih.gov/pubmed/22961589.

41. Shao, J., S. W. Tanner, N. Thompson, and T. E. Cheatham. 2007. Clustering Molecular Dynamics Trajectories: 1. Characterizing the Performance of Different Clustering Algorithms. J Chem Theory Comput. 3(6):2312–2334, doi: 10.1021/ct700119m, https://www.ncbi.nlm.nih.gov/pubmed/26636222.

42. Richardson, J. S., B. Schneider, L. W. Murray, G. J. Kapral, R. M. Immormino, J. J. Headd, D. C. Richardson, D. Ham, E. Hershkovits, L. D. Williams, K. S. Keating, A. M. Pyle, D. Micallef, J. Westbrook, and H. M. Berman. 2008. RNA backbone: consensus all-angle conformers and modular string nomenclature (an RNA Ontology Consortium contribution). RNA. 14(3):465–481, doi: 10.1261/rna.657708, http://www.ncbi.nlm.nih.gov/pubmed/18192612.

43. Cavanagh, J., W. J. Fairbrother, A. G. Palmer, and N. J. Skelton. 1996. Protein NMR Spectroscopy: Principles and Practice Academic Press, San Diego.

44. Reiter, N. J., H. Blad, F. Abildgaard, and S. E. Butcher. 2004. Dynamics in the U6 RNA intramolecular stem-loop: a base flipping conformational change. Biochemistry. 43(43):13739–13747, doi: 10.1021/bi048815y, http://www.ncbi.nlm.nih.gov/pubmed/15504036.

45. Macura, S., W. M. Westler, and J. L. Markley. 1994. Two-dimensional exchange spectroscopy of proteins. Methods Enzymol. 239:106–144, doi: S0076-6879(94)39005-3 [pii], http://www.ncbi.nlm.nih.gov/pubmed/7830582.

46. Yildirim, I., H. A. Stern, S. D. Kennedy, J. D. Tubbs, and D. H. Turner. 2010. Reparameterization of RNA chi Torsion Parameters for the AMBER Force Field and Comparison to NMR Spectra for Cytidine and Uridine. J Chem Theory Comput. 6(5):1520–1531, doi: 10.1021/ct900604a, http://www.ncbi.nlm.nih.gov/pubmed/20463845.

47. Yildirim, I., S. D. Kennedy, H. A. Stern, J. M. Hart, R. Kierzek, and D. H. Turner. 2012. Revision of AMBER Torsional Parameters for RNA Improves Free Energy Predictions for Tetramer Duplexes with GC and iGiC Base Pairs. J Chem Theory Comput. 8(1):172–181, doi: 10.1021/ct200557r, http://www.ncbi.nlm.nih.gov/pubmed/22249447.

48. Chen, A. A., and A. E. Garcia. 2013. High-resolution reversible folding of hyperstable RNA tetraloops using molecular dynamics simulations. Proc Natl Acad Sci U S A. 110(42):16820–16825, doi: 10.1073/pnas.1309392110, https://www.ncbi.nlm.nih.gov/pubmed/24043821.

49. Aytenfisu, A. H., A. Spasic, A. Grossfield, H. A. Stern, and D. H. Mathews. 2017. Revised RNA Dihedral Parameters for the Amber Force Field Improve RNA Molecular Dynamics. J Chem Theory Comput. 13(2):900–915, doi: 10.1021/acs.jctc.6b00870, http://www.ncbi.nlm.nih.gov/pubmed/28048939.

50. Tan, D., S. Piana, R. M. Dirks, and D. E. Shaw. 2018. RNA force field with accuracy comparable to state-of-the-art protein force fields. Proc Natl Acad Sci U S A. 115(7):E1346–E1355, doi: 10.1073/pnas.1713027115, https://www.ncbi.nlm.nih.gov/pubmed/29378935.

51. Denning, E. J., U. D. Priyakumar, L. Nilsson, and A. D. Mackerell, Jr. 2011. Impact of 2’- hydroxyl sampling on the conformational properties of RNA: update of the CHARMM all-atom additive force field for RNA. Journal of Computational Chemistry. 32(9):1929–1943, doi: 10.1002/jcc.21777, http://www.ncbi.nlm.nih.gov/pubmed/21469161.

52. Frohlking, T., V. Mlynsky, M. Janecek, P. Kuhrova, M. Krepl, P. Banas, J. Sponer, and G. Bussi. 2022. Automatic Learning of Hydrogen-Bond Fixes in the AMBER RNA Force Field. J Chem Theory Comput. 18(7):4490–4502, doi: 10.1021/acs.jctc.2c00200, https://www.ncbi.nlm.nih.gov/pubmed/35699952.

53. Ananth, P., G. Goldsmith, and N. Yathindra. 2013. An innate twist between Crick’s wobble and Watson-Crick base pairs. RNA. 19(8):1038–1053, doi: 10.1261/rna.036905.112, https://www.ncbi.nlm.nih.gov/pubmed/23861536.

54. Masquida, B., and E. Westhof. 2000. On the wobble GoU and related pairs. RNA. 6(1):9–15, doi: 10.1017/s1355838200992082, https://www.ncbi.nlm.nih.gov/pubmed/10668794.

55. Mizuno, H., and M. Sundaralingam. 1978. Stacking of Crick Wobble pair and Watson-Crick pair: stability rules of G-U pairs at ends of helical stems in tRNAs and the relation to codon-anticodon Wobble interaction. Nucleic Acids Res. 5(11):4451–4461, doi: 10.1093/nar/5.11.4451, https://www.ncbi.nlm.nih.gov/pubmed/724522.

56. Bu, F., Y. Adam, R. W. Adamiak, M. Antczak, B. R. H. de Aquino, N. G. Badepally, R. T. Batey, E. F. Baulin, P. Boinski, M. J. Boniecki, J. M. Bujnicki, K. A. Carpenter, J. Chacon, S. J. Chen, W. Chiu, P. Cordero, N. K. Das, R. Das, W. K. Dawson, F. DiMaio, F. Ding, A. C. Dock-Bregeon, N. V. Dokholyan, R. O. Dror, S. Dunin-Horkawicz, S. Eismann, E. Ennifar, R. Esmaeeli, M. A. Farsani, A. R. Ferre-D’Amare, C. Geniesse, G. E. Ghanim, H. V. Guzman, I. V. Hood, L. Huang, D. S. Jain, F. Jaryani, L. Jin, A. Joshi, M. Karelina, J. S. Kieft, W. Kladwang, S. Kmiecik, D. Koirala, M. Kollmann, R. C. Kretsch, M. Kurcinski, J. Li, S. Li, M. Magnus, B. Masquida, S. N. Moafinejad, A. Mondal, S. Mukherjee, T. H. D. Nguyen, G. Nikolaev, C. Nithin, G. Nye, I. P. N. Pandaranadar Jeyeram, A. Perez, P. Pham, J. A. Piccirilli, S. P. Pilla, R. Pluta, S. Poblete, A. Ponce-Salvatierra, M. Popenda, L. Popenda, F. Pucci, R. Rangan, A. Ray, A. Ren, J. Sarzynska, C. M. Sha, F. Stefaniak, Z. Su, K. C. Suddala, M. Szachniuk, R. Townshend, R. J. Trachman, 3rd, J. Wang, W. Wang, A. Watkins, T. K. Wirecki, Y. Xiao, P. Xiong, Y. Xiong, J. Yang, J. D. Yesselman, J. Zhang, Y. Zhang, Z. Zhang, Y. Zhou, T. Zok, D. Zhang, S. Zhang, A. Zyla, E. Westhof, and Z. Miao. 2025. RNA-Puzzles Round V: blind predictions of 23 RNA structures. Nat Methods. 22(2):399-411, doi: 10.1038/s41592-024-02543-9, https://www.ncbi.nlm.nih.gov/pubmed/39623050.

57. Leontis, N. B., J. Stombaugh, and E. Westhof. 2002. The non-Watson-Crick base pairs and their associated isostericity matrices. Nucleic Acids Res. 30(16):3497–3531, doi: 10.1093/nar/gkf481, http://www.ncbi.nlm.nih.gov/pubmed/12177293.

