## Supplementary Figures and Tables for "NMR and molecular dynamics demonstrate the RNA internal loop GAGU is dynamic and adjacent base pairs determine conformational preference"

**Figure S1**. NMR spectra of the imino proton region of the duplexes GgaguC, AgaguU, and UgaguA. Peaks are labeled with I (conformation I) or 'a' (conformations II/III). Dilution by 100 fold identifies peaks that arise from minor populations of hairpin (indicated by asterisk). Hairpin peaks become relatively larger compared to duplex peaks when the concentration of RNA is sufficiently reduced. The dilution of the duplex CgaguG can be found in (1) . The inset graph with AgaguU shows the intensity as a function of mixing time of G2a on the diagonal of 2D NOESY spectra (black) and the intensity of the G2a-G2I exchange cross-peak (blue; intensity multiplied by 10) demonstrating that conformation I and the alternate conformation interconvert on the order of 25-50 ms.

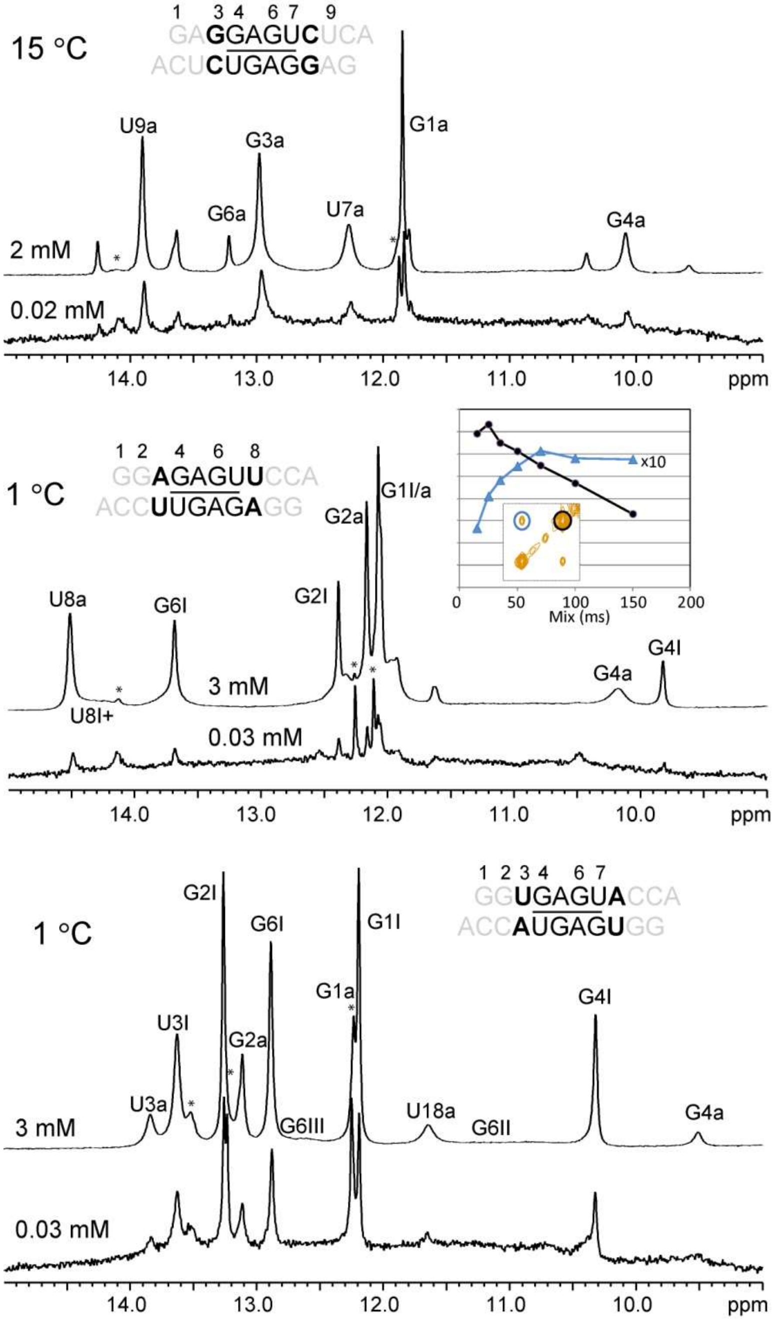

**Figure S2.** 2D NOEs of GgaguC and of related AG 2×2 internal loops showing pattern of contacts between GH8 and sugar protons typical when alpha and gamma torsions are in the “crankshaft” conformation (1c): the H8-H5’’ distance is shorter than the H8-H5’ distance. The distances are approximately equal in A-form (1a). Confirmation of the gamma torsion in the AG 2×2 loops was made by measures of H4’-H5’/’’ scalar couplings (1). This coupling could not be resolved in GgaguC. Vertical lines are G6H8.

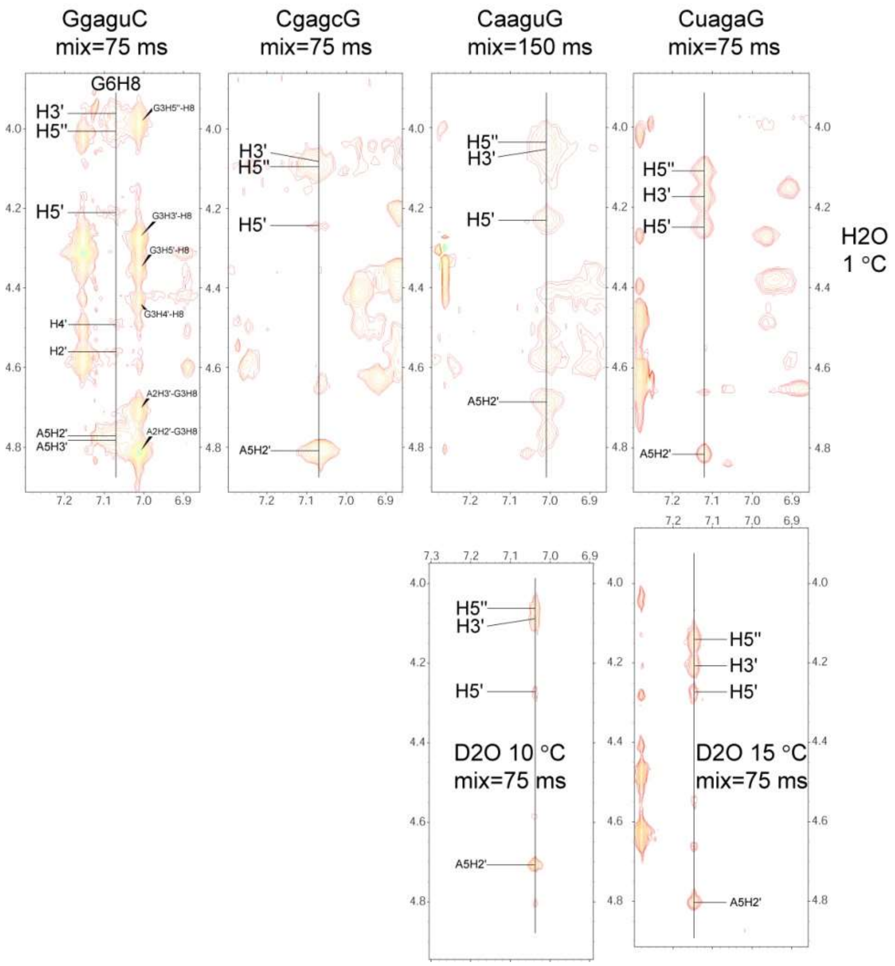

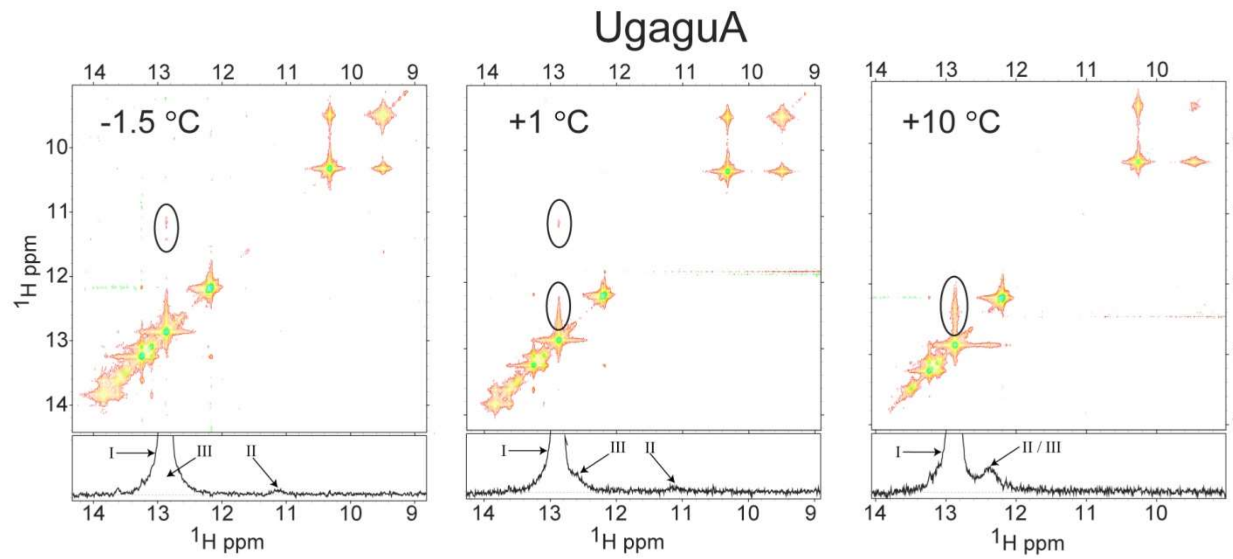


**Figure S3**: Identification of separate G6H1 peaks for conformations II and III in UgaguA at -1.5 °C and 1 °C but only one peak at 10 °C. The 1D spectral slice below each 2D-NOESY spectrum is taken horizontally at the position of G6H1 in conformation I. Exchange cross-peaks from conformation I to both conformation II and III are observed (solid ovals in 2D; arrows in 1D).

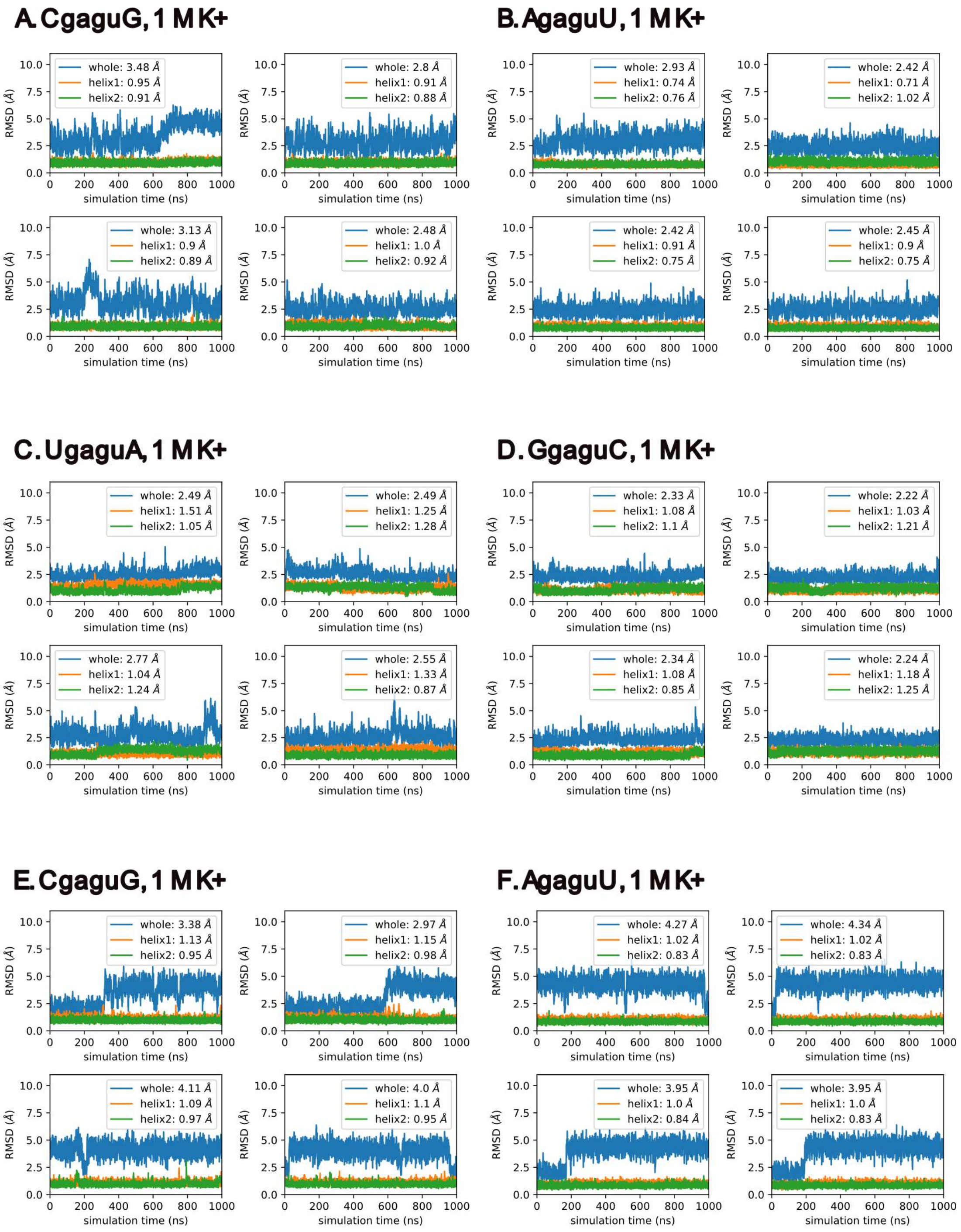

A. CgaguG, 1 M K+
whole: 3.48 Å
helix1: 0.95 Å
helix2: 0.91 Å
whole: 2.8 Å
helix1: 0.91 Å
helix2: 0.88 Å
whole: 3.13 Å
helix1: 0.9 Å
helix2: 0.89 Å
whole: 2.48 Å
helix1: 1.0 Å
helix2: 0.92 Å
B. AgaguU, 1 M K+
whole: 2.93 Å
helix1: 0.74 Å
helix2: 0.76 Å
whole: 2.42 Å
helix1: 0.71 Å
helix2: 1.02 Å
whole: 2.42 Å
helix1: 0.91 Å
helix2: 0.75 Å
whole: 2.45 Å
helix1: 0.9 Å
helix2: 0.75 Å
C. UgaguA, 1 M K+
whole: 2.49 Å
helix1: 1.51 Å
helix2: 1.05 Å
whole: 2.49 Å
helix1: 1.25 Å
helix2: 1.28 Å
whole: 2.77 Å
helix1: 1.04 Å
helix2: 1.24 Å
whole: 2.55 Å
helix1: 1.33 Å
helix2: 0.87 Å
D. GgaguC, 1 M K+
whole: 2.33 Å
helix1: 1.08 Å
helix2: 1.1 Å
whole: 2.22 Å
helix1: 1.03 Å
helix2: 1.21 Å
whole: 2.34 Å
helix1: 1.08 Å
helix2: 0.85 Å
whole: 2.24 Å
helix1: 1.18 Å
helix2: 1.25 Å
E. CgaguG, 1 M K+
whole: 3.38 Å
helix1: 1.13 Å
helix2: 0.95 Å
whole: 2.97 Å
helix1: 1.15 Å
helix2: 0.98 Å
whole: 4.11 Å
helix1: 1.09 Å
helix2: 0.97 Å
whole: 4.0 Å
helix1: 1.1 Å
helix2: 0.95 Å
F. AgaguU, 1 M K+
whole: 4.27 Å
helix1: 1.02 Å
helix2: 0.83 Å
whole: 4.34 Å
helix1: 1.02 Å
helix2: 0.83 Å
whole: 3.95 Å
helix1: 1.0 Å
helix2: 0.84 Å
whole: 3.95 Å
helix1: 1.0 Å
helix2: 0.83 Å
RMSD (Å)
simulation time (ns)

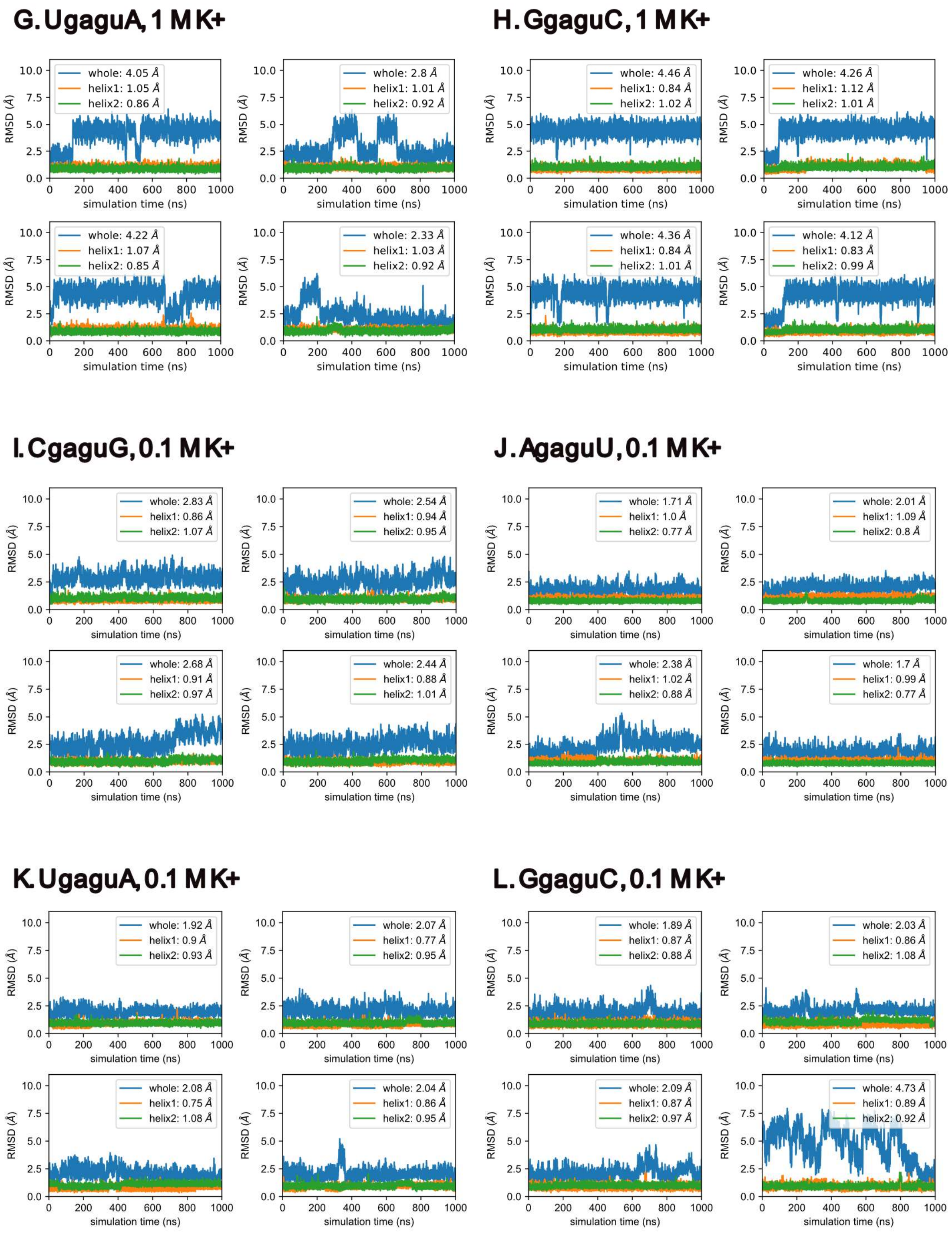

G. UgaguA, 1 M K+
whole: 4.05 Å
helix1: 1.05 Å
helix2: 0.86 Å
whole: 2.8 Å
helix1: 1.01 Å
helix2: 0.92 Å
whole: 4.22 Å
helix1: 1.07 Å
helix2: 0.85 Å
whole: 2.33 Å
helix1: 1.03 Å
helix2: 0.92 Å
H. GgaguC, 1 M K+
whole: 4.46 Å
helix1: 0.84 Å
helix2: 1.02 Å
whole: 4.26 Å
helix1: 1.12 Å
helix2: 1.01 Å
whole: 4.36 Å
helix1: 0.84 Å
helix2: 1.01 Å
whole: 4.12 Å
helix1: 0.83 Å
helix2: 0.99 Å
I. CgaguG, 0.1 M K+
whole: 2.83 Å
helix1: 0.86 Å
helix2: 1.07 Å
whole: 2.54 Å
helix1: 0.94 Å
helix2: 0.95 Å
whole: 2.68 Å
helix1: 0.91 Å
helix2: 0.97 Å
whole: 2.44 Å
helix1: 0.88 Å
helix2: 1.01 Å
J. AgaguU, 0.1 M K+
whole: 1.71 Å
helix1: 1.0 Å
helix2: 0.77 Å
whole: 2.01 Å
helix1: 1.09 Å
helix2: 0.8 Å
whole: 2.38 Å
helix1: 1.02 Å
helix2: 0.88 Å
whole: 1.7 Å
helix1: 0.99 Å
helix2: 0.77 Å
K. UgaguA, 0.1 M K+
whole: 1.92 Å
helix1: 0.9 Å
helix2: 0.93 Å
whole: 2.07 Å
helix1: 0.77 Å
helix2: 0.95 Å
whole: 2.08 Å
helix1: 0.75 Å
helix2: 1.08 Å
whole: 2.04 Å
helix1: 0.86 Å
helix2: 0.95 Å
L. GgaguC, 0.1 M K+
whole: 1.89 Å
helix1: 0.87 Å
helix2: 0.88 Å
whole: 2.03 Å
helix1: 0.86 Å
helix2: 1.08 Å
whole: 2.09 Å
helix1: 0.87 Å
helix2: 0.97 Å
whole: 4.73 Å
helix1: 0.89 Å
helix2: 0.92 Å
RMSD (Å)
simulation time (ns)

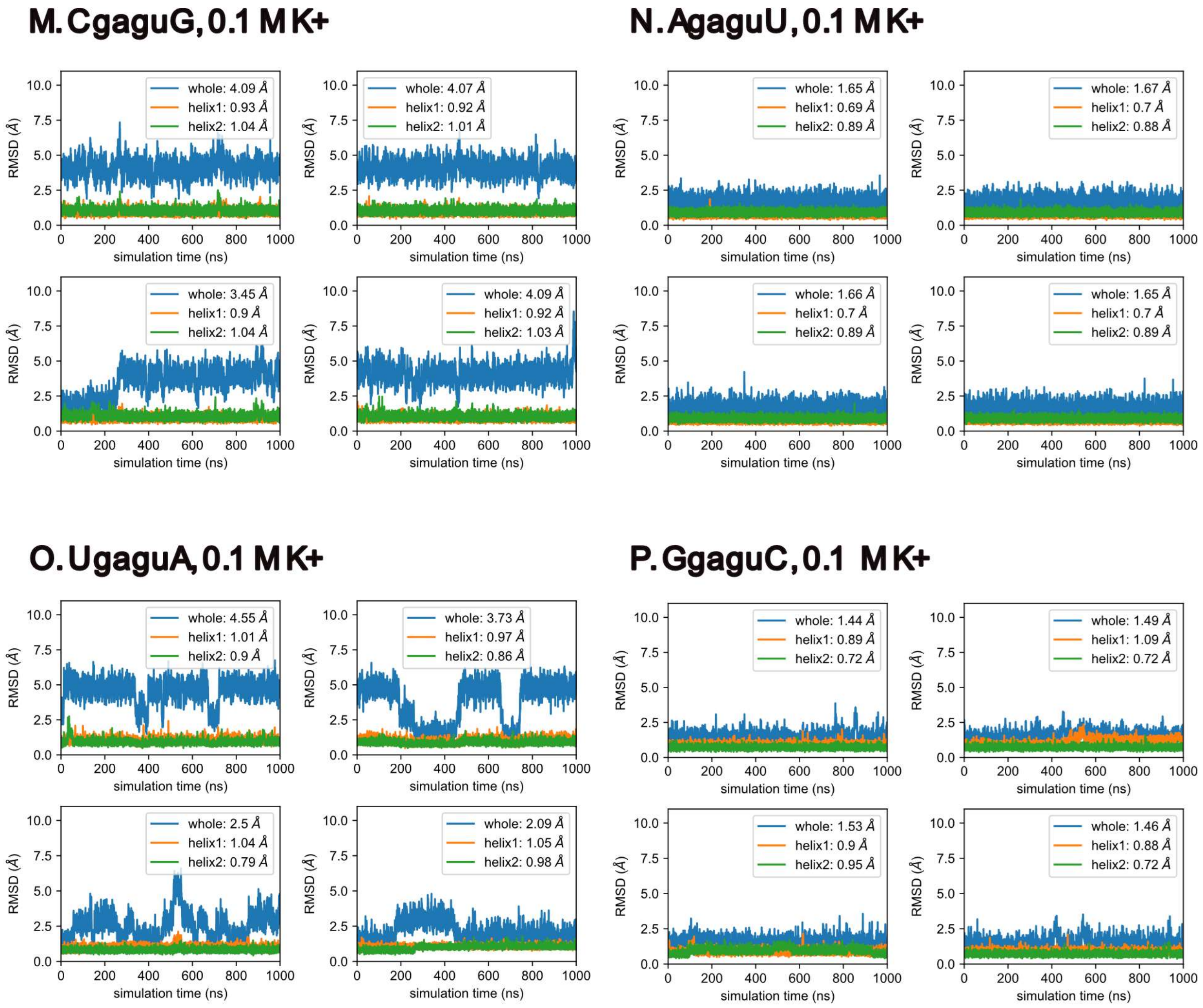


**Figure S4.** Heavy atom RMSD of trajectory structures relative to the energy-minimized structure for CgaguG, AgaguU, UgaguA and GgaguC. Panels A, B, C, D refer to simulations starting in conformation I in 1 M $K^+$. Panels E, F, G, H refer to simulations starting in conformation III in 1 M $K^+$. Panels I, J, K, L refer to simulations starting in conformation I in 0.1 M $K^+$. Panels M, N, O, P refer to simulations starting in conformation III in 0.1 M $K^+$. Each panel contains four subplots: upper left (replicate 1), upper right (replicate 2), lower left (replicate 3) and lower right (replicate 4).

| **Starting Conformation III** | | | | |
|---|---|---|---|---|
| **Sequence** | **G4-U7*** | **G4*-U7** | **A5-G6*** | **A5*-G6** |
| UgaguA | 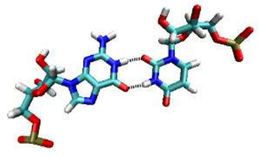 | 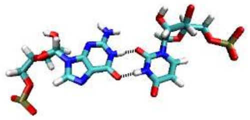 | 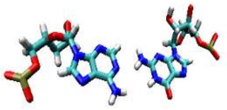 | 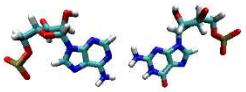 |
| **Starting Conformation I** | | | | |
| **Sequence** | **G4-G6*** | **G4*-G6** | **A5:A5*** | **-** |
| GgaguC | 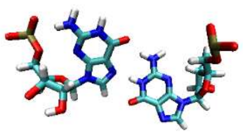 | 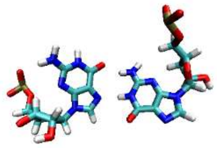 | 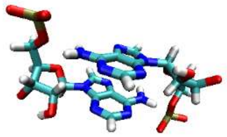 | |

Figure S5: Snapshots of cluster centroids of intermediate structures (Table 5) present in simulations in 0.1 M $K^+$.

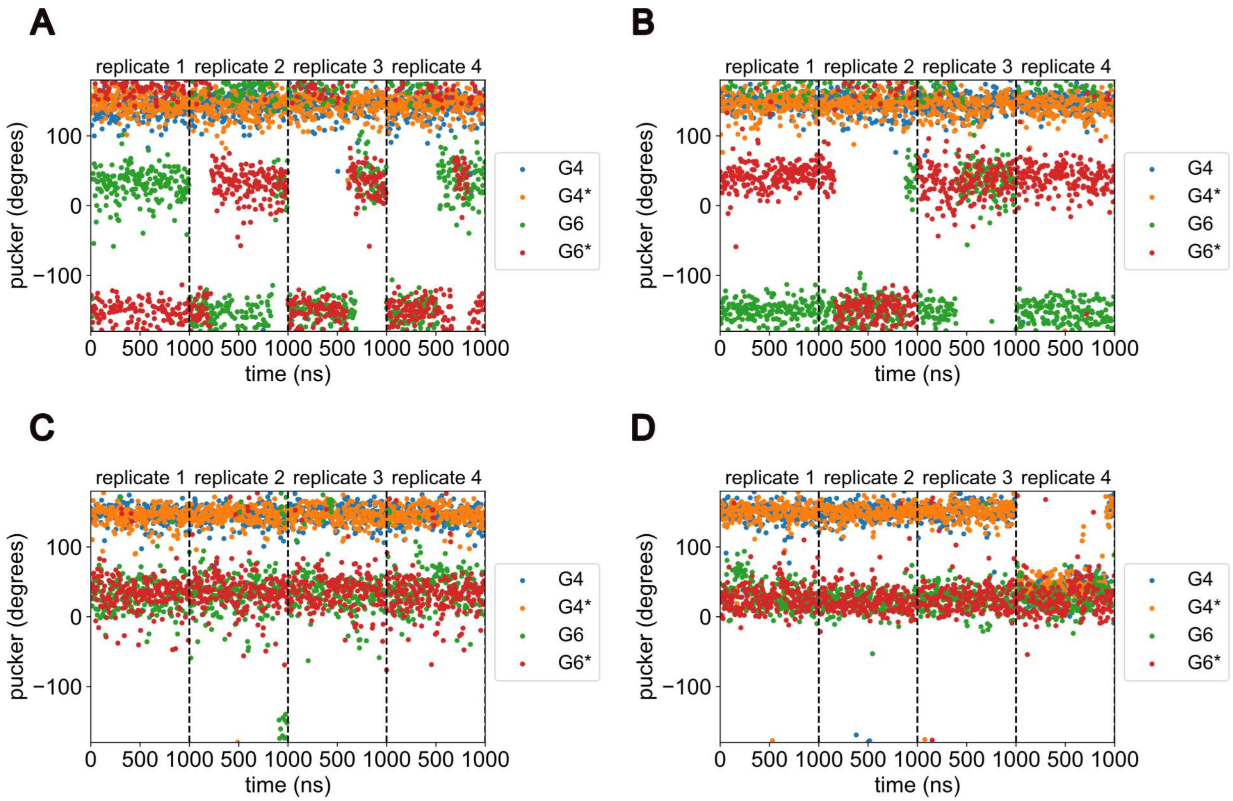


**Figure S6**: Time plot of sugar pucker of structures generated in MD simulations of CgaguG (panel A), AgaguU (panel B), UgaguA (panel C) and GgaguC (panel D) starting in conformation I in 0.1 M $K^+$, for loop closing pairs G4/G4* and G6/G6*. Original trajectory (200 ps / frame) was downsampled to 1 in every 50 frames.

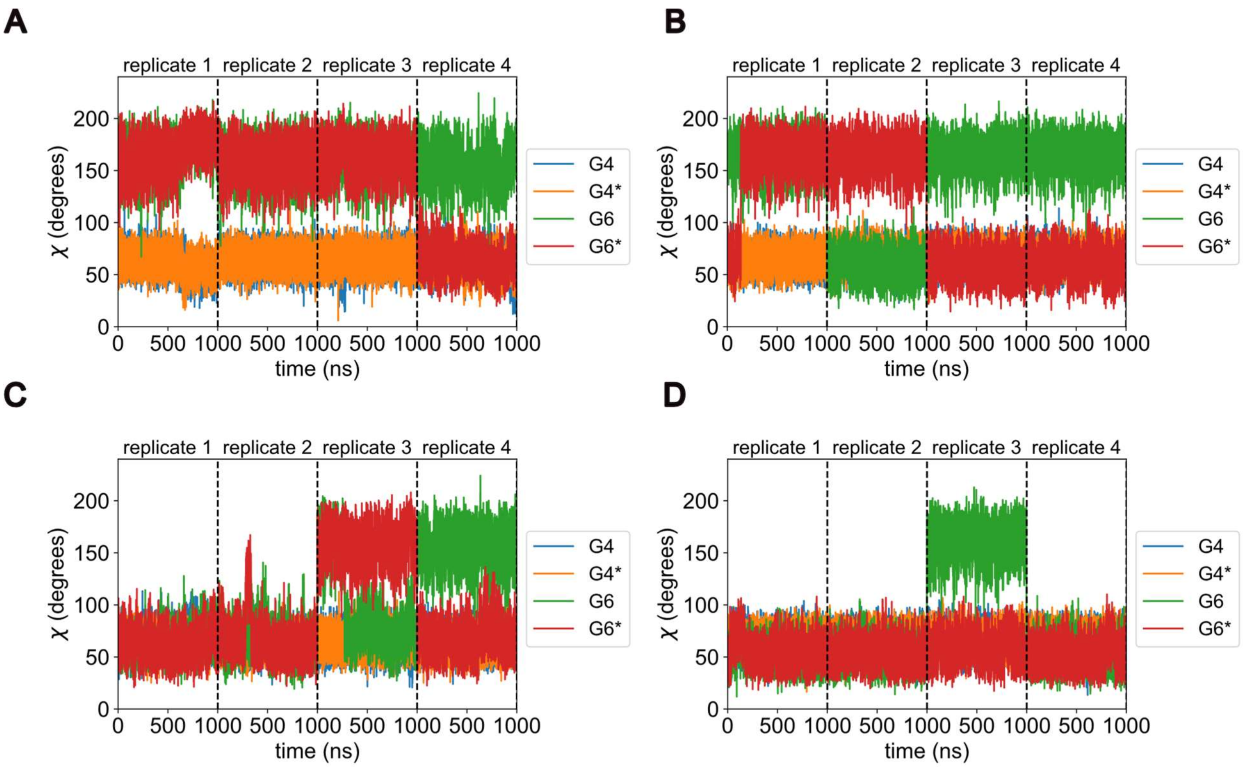


Figure S7: Time series plots of χ dihedral angles for G4, G4*, G6 and G6* for CgaguG (Panel A), AgaguU (Panel B), UgaguA (Panel C) and GgaguC (Panel D) simulations in 1.0 M $K^+$ starting in conformation I.

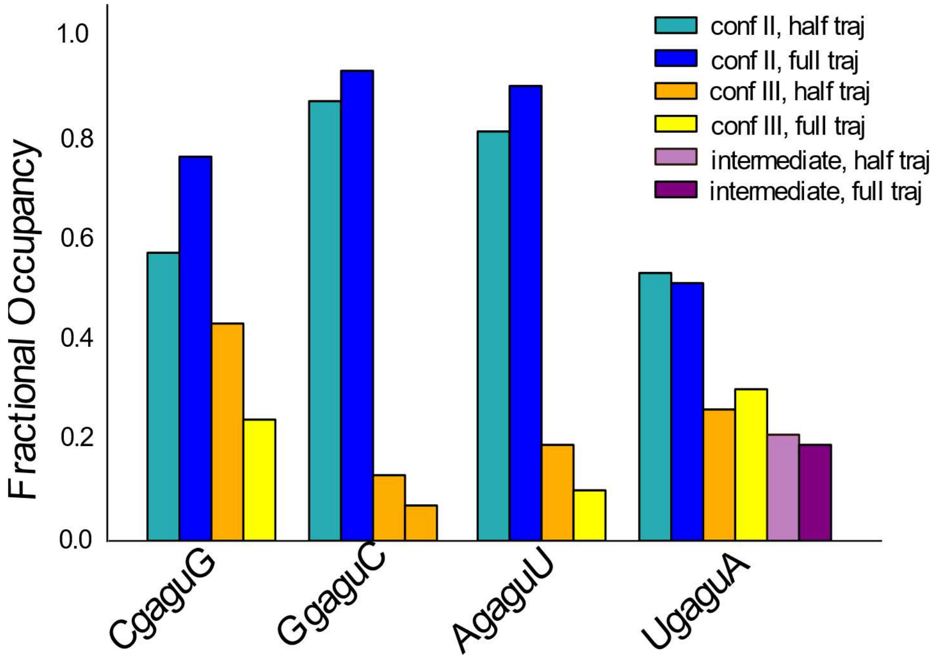


**Figure S8**. Simulation in 1 M $K^+$. The average fractional population of clusters of conformation II/III structures over four replicates in the first 500 ns ("half traj") and the entire 1 μs ("full traj") of simulation. Results show that extending the simulation from 500 ns to 1 μs does not substantially alter cluster statistics. Duplex UgaguA has a third cluster of intermediate structures with II-like GU and III-like AG pairs.

A

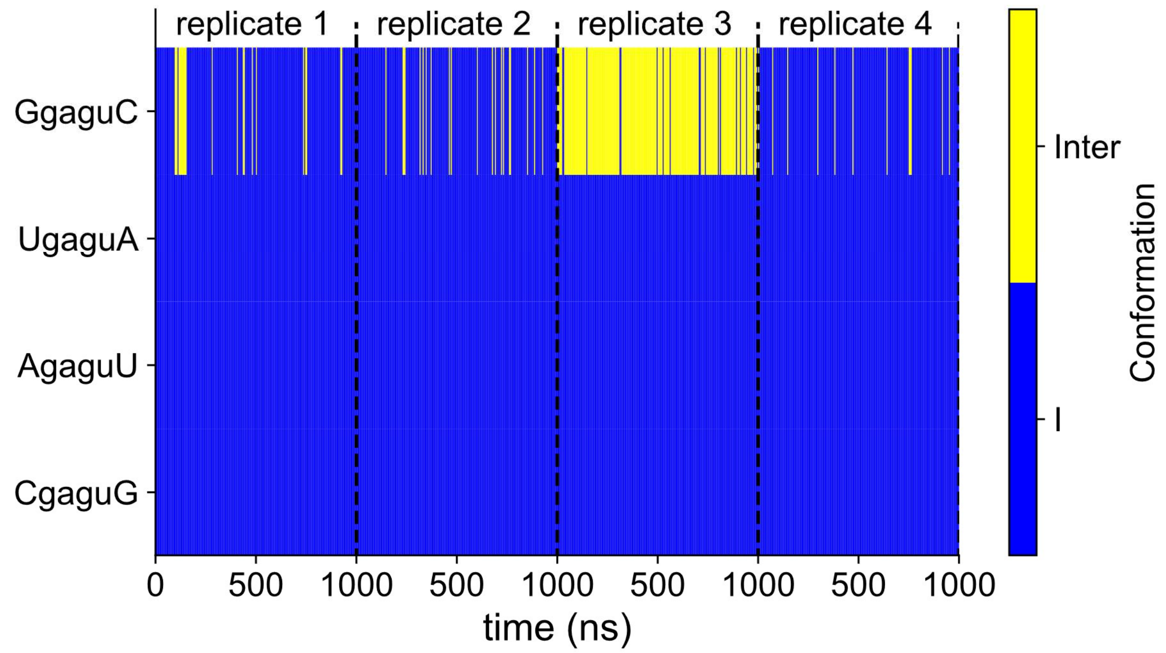


B

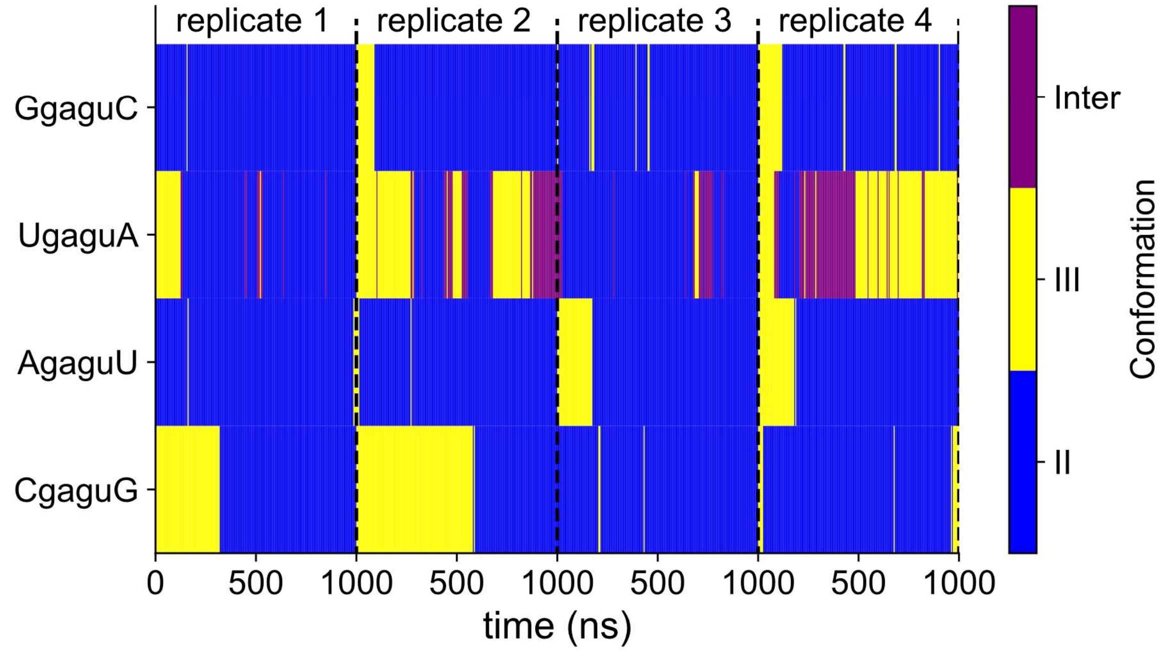


Fig S9: Time series plot of cluster index for structures generated in MD simulation in 1 M $K^+$ solvent. Simulations starting in conformation I (panel A) showed clear signs of lack of convergence marked by the presence of intermediate structures in replicate three of GgaguC. Simulations starting in conformation III (panel B) had structures in conformation as the dominant population for all duplexes, with UgaguA having a substantial population of structures in intermediate state.

**Table S1.** Chemical shifts of all assigned proton resonances of conformations I and II/III at 1 °C in four duplexes containing 5'GAGU/3'UGAG.

| Sequence | GgaguC | | | UgaguA | | | AgaguU | | | CgaguG | | |
|---|---|---|---|---|---|---|---|---|---|---|---|---|
| Conformation | | III | I | | II / III | I | | II / III | I | | II | I |
| H1 | **G1** | 11.81 | 11.75 | **G1** | 12.23 | 12.19 | **G1** | 12.07 | 12.07 | **G1** | 12.01 | 11.89 |
| H8 | | 8.07 | | | 8.09 | 8.09 | | 8.08 | 8.08 | | | 8.07 |
| H1' | | 5.81 | | | 5.88 | 5.88 | | 5.88 | 5.88 | | | 5.81 |
| H2' | | 4.78 | | | 4.91 | 4.91 | | | | | | 4.83 |
| H3' | | 4.58 | | | | | | | | | | |
| H4' | | 4.39 | | | | | | | | | | |
| H5' | | 4.07 | | | | | | | | | | |
| H5'' | | 3.92 | | | | | | | | | | |
| amino1 | | 6.12 | | | 6.23 | 6.23 | | 6.16 | | | | |
| amino2 | | 7.71 | | | 8.07 | 8.07 | | 8.00 | | | | |
| H1 | **A2** | | | **G2** | 13.11 | 13.26 | **G2** | 12.16 | 12.39 | **A2** | | |
| H2 | | 7.43 | 7.19 | | | | | | | | 7.71 | 7.54 |
| H8 | | 7.95 | | | 7.42 | 7.29 | | 7.39 | 7.24 | | | 7.90 |
| H1' | | 5.96 | | | 5.79 | 5.73 | | 5.77 | 5.69 | | 5.96 | 5.90 |
| H2' | | 4.81 | | | | 4.48 | | | 4.53 | | | |
| H3' | | 4.71 | | | | | | | | | | |
| H4' | | 4.49 | | | | | | | | | | |
| H5' | | 4.60 | | | | | | | | | | |
| H5'' | | 4.16 | | | | | | | | | | |
| amino1 | | 7.76 | | | | 5.88 | | | | | | |
| amino2 | | 6.32 | | | | 8.36 | | 8.00 | 7.88 | | | |
| H1/H3 | **G3** | 13.02 | 13.22 | **U3** | 13.85 | 13.63 | **A3** | | | **C3** | | |
| H2/H5 | | | | | | 4.91 | | 7.24 | 7.43 | | | 4.94 |
| H6/H8 | | 7.01 | | | 7.60 | 7.42 | | 7.63 | 7.44 | | 7.32 | 7.12 |
| H1' | | 5.46 | | | 5.48 | 5.24 | | 5.91 | 5.64 | | 5.32 | 5.01 |
| H2' | | 4.32 | | | | | | 4.56 | 4.16 | | | |
| H3' | | 4.28 | | | | 4.37 | | | 4.53 | | | |
| H4' | | 4.44 | | | | | | | | | | |
| H5' | | 4.36 | | | | | | | | | | |
| H5'' | | 3.99 | | | | | | | | | | |
| amino1 | | 5.79 | | | | | | | | | 7.95 | 8.12 |
| amino2 | | 7.57 | | | | | | | | | 6.78 | 6.79 |
| C2 | | | | | | | | 152.70 | 154.40 | | | |
| H1 | **G4** | 10.14 | 9.68 | **G4** | 9.50 | 10.32 | **G4** | 10.18 | 9.82 | **G4** | 9.37 | 10.18 |
| H8 | | 7.15 | 7.66 | | | 7.60 | | 6.93 | 7.52 | | 7.30 | 7.58 |
| H1' | | 5.60 | | | 5.60 | 5.66 | | | 5.57 | | 5.56 | 5.62 |
| H2' | | 4.27 | | | | | | | | | | |

| | | | | | | | | | | | | |
|---|---|---|---|---|---|---|---|---|---|---|---|---|
| H3' | | 4.58 | | | | | | | | | | |
| H4' | | 4.36 | | | | | | | | | | |
| H5' | | 4.50 | | | | | | | | | | |
| H5'' | | 4.02 | | | | | | | | | | |
| amino | | 6.21 | | | 5.91 | | | 6.21 | | | 5.44 | |
| H2 | **A5** | 7.89 | 7.70 | **A5** | 7.75 | 7.69 | **A5** | 7.87 | 7.68 | **A5** | 7.91 | 7.68 |
| H8 | | 8.16 | 7.55 | | 8.08 | 7.60 | | 8.00 | 7.57 | | 7.69 | 7.58 |
| H1' | | 6.01 | 5.70 | | 5.89 | 5.69 | | 5.90 | 5.65 | | 5.63 | 5.69 |
| H2' | | 4.78 | | | | | | 4.72 | 3.96 | | | 4.00 |
| H3' | | 4.78 | | | | | | | | | | |
| H4' | | 4.43 | | | | | | | | | | |
| H5' | | | | | | 2.77 | | | 2.92 | | | 2.74 |
| H5'' | | | | | | 3.59 | | | 3.59 | | | 3.60 |
| amino | | 7.65 | | | | | | | | | | |
| C2 | | | | | 153.90 | 153.50 | | | 153.60 | | | |
| | | | | **G6** | | | | | | | | |
| H1 | **G6** | 13.06 | 13.71 | **II/I** | 11.13 | 12.88 | **G6** | 12.35 | 13.68 | **G6** | 10.99 | 13.31 |
| | | | | **III** | 12.64 | | | | | | | |
| H8 | | 7.07 | 7.79 | | 7.16 | 7.71 | | 7.23 | 7.84 | | 7.51 | 7.78 |
| H1' | | 5.48 | 5.84 | | 5.40 | 5.80 | | 5.43 | 5.83 | | 5.39 | 5.84 |
| H2' | | 4.56 | | | | 5.51 | | ? | | | | 5.65 |
| H3' | | 3.96 | | | | 4.89 | | ? | | | | 4.84 |
| H4' | | 4.49 | | | | | | | | | | |
| H5' | | 4.22 | | | | | | | | | | |
| H5'' | | 4.01 | | | | | | ? | | | | |
| amino | | 6.07 | 6.96 | | 6.08 | 6.68 | | 6.25 | 6.90 | | 6.25 | 6.59 |
| H3 | **U7** | 12.32 | | **U7** | 11.63 | | **U7** | ? | | **U7** | | |
| H5 | | 5.18 | 5.95 | | 5.21 | 5.98 | | 5.17 | 5.97 | | 5.21 | 5.96 |
| H6 | | 7.60 | 7.93 | | 7.56 | 8.04 | | 7.60 | 7.99 | | 7.53 | 8.04 |
| H1' | | 5.57 | 6.09 | | 5.56 | 6.18 | | 5.61 | 6.16 | | 5.54 | 6.15 |
| H2' | | 3.99 | | | 4.12 | | | ? | | | | |
| H3' | | 4.49 | | | | | | | | | | |
| H4' | | 4.36 | | | | | | | | | | |
| H1/H3 | **C8** | | | **A8** | | | **U8** | 14.51 | 14.24 | **G8** | 12.84 | 12.95 |
| H2/H5 | | 5.57 | 5.93 | | 7.16 | 7.46 | | 5.57 | 5.83 | | | |
| H6/H8 | | 8.03 | 7.81 | | 8.35 | 8.17 | | 8.11 | 7.96 | | 8.06 | 7.76 |
| H1' | | 5.49 | | | 5.83 | 5.27 | | 5.53 | 4.80 | | 5.76 | 5.02 |
| H2' | | 4.12 | | | | 4.42 | | 4.38 | | | | |
| H3' | | 4.49 | | | | | | ? | | | | |
| H4' | | 4.32 | | | | | | | | | | |
| amino1 | | 8.34 | 8.75 | | | | | | | | | |
| amino2 | | 7.07 | 6.97 | | | | | | | | | |

| | | | | | | | | | | | | |
|---|---|---|---|---|---|---|---|---|---|---|---|---|
| C2 | | | | | 153.10 | 153.90 | | | | | | |
| H3 | **U9** | 13.91 | 14.30 | **C9** | | | **C9** | | | **U9** | 14.29 | 14.51 |
| H5 | | 5.32 | | | 5.15 | 5.19 | | 5.57 | 5.69 | | | 5.06 |
| H6 | | 7.81 | | | 7.50 | 7.56 | | 7.83 | 7.91 | | 7.73 | 7.80 |
| H1' | | 5.38 | | | | 5.33 | | 5.50 | 5.49 | | | 5.41 |
| H2' | | 4.42 | | | | 4.12 | | | | | | 4.38 |
| H3' | | 4.42 | | | | | | | | | | |
| H4' | | 4.34 | | | | | | | | | | |
| amino1 | | | | | 8.31 | 8.31 | | 8.27 | 8.37 | | | |
| amino2 | | | | | 6.87 | 6.89 | | 6.91 | 7.05 | | | |
| H5 | **C10** | 5.53 | | **C10** | 5.34 | 5.34 | **C10** | 5.39 | 5.39 | **C10** | | 5.52 |
| H6 | | 7.74 | | | 7.52 | 7.52 | | 7.59 | 7.56 | | | 7.65 |
| H1' | | 5.48 | | | 5.37 | 5.37 | | 5.37 | 5.32 | | | 5.44 |
| H2' | | 4.33 | | | | | | | | | | 4.35 |
| H3' | | 4.42 | | | | | | | | | | |
| H4' | | 4.35 | | | | | | | | | | |
| amino1 | | 8.02 | 8.00 | | 8.15 | 8.15 | | 8.11 | 8.11 | | 8.22 | 8.11 |
| amino2 | | 6.98 | | | 6.88 | 6.88 | | 6.93 | 6.93 | | 6.98 | 6.97 |
| H2 | **A11** | 7.12 | | **A11** | 7.26 | 7.27 | **A11** | 7.19 | 7.22 | **A11** | | 7.24 |
| H8 | | 7.92 | | | 7.94 | 7.94 | | 7.93 | 7.93 | | | 7.96 |
| H1' | | 5.84 | | | 5.92 | 5.92 | | 5.89 | 5.89 | | | 5.87 |
| H2' | | 3.95 | | | | | | 3.98 | 3.98 | | | |
| H3' | | 4.22 | | | | | | 4.25 | | | | |
| H4' | | 4.15 | | | | | | | | | | |
| C2 | | | | | | 154.00 | | 154.00 | | | | |

**Table S2**. G6/G4 H1'-H8 1/$r^6$ ratio for simulations starting in conformation I for CgaguG, AgaguU, UgaguA and GgaguC in both 0.1 M $K^+$ and 1.0 $K^+$ solvent. Values are mean and standard deviation of the ratio for the four replicates. Also shown are the NMR NOE ratios for G6 and G4 H1'-H8. Values are mean and standard deviation from two NOESY spectra and four integration boxes. The low population of conformation I prevents determination of this ratio in GgaguC.

---

| G6/G4 H1'-H8 1/$r^6$ Ratio | | | |
|---|---|---|---|
| Duplex | 0.1 M $K^+$ Simulations | 1.0 M $K^+$ Simulations | NMR (G6/G4 H1'-H8 NOE Ratio) |
| CgaguG | 0.54 ± 0.07 | 0.31 ± 0.20 | 0.83 ± 0.01 |
| AgaguU | 0.59 ± 0.20 | 0.51 ±0.16 | 0.90 ± 0.02 |
| UgaguA | 1.03 ± 0.02 | 0.80 ± 0.25 | 0.60 ± 0.04 |
| GgaguC | 1.03 ± 0.01 | 0.94 ± 0.19 | NA |

**Table S3.**
NMR structure statistics for the duplex 5'GAGGAGUCUCA/3'ACUCUGAGGAG (GgaguC).

| | |
|---|---|
| Distance restraints | |
| Total NOEs | 173 |
| Intranucleotide | 72 |
| Internucleotide | 101 |
| Long range (\|i-j\|≥2) | 32 |
| Violations > 0.2 Å | 0 |
| Hydrogen bond restraints | 24 |
| Dihedral angle restraints | 129s |
| Violations > 5° | 30 |
| RMSD to restraints | |
| NOEs | .0001 Å |
| Dihedral angles | 0.41° |
| Energies (kcal/mole) | |
| Total (system) | |
| NOE | |
| Dihedral | |
| Structures calculated/accepted | 600/30 |
| RMSD to centroid structure (Å) | |
| Heavy atoms | 0.86 |
| All atoms | 0.88 |

**Table S4**: Dihedal angle of the centroid structure of GgaguC starting in conformation III for 0.1 M K+ simulations.

| Base | α | β | γ | Δ |
|---|---|---|---|---|
| G6 | -68.80 | 171.45 | 64.10 | 84.70 |
| G6* | -73.80 | 168.09 | 64.04 | 76.65 |
| Mean (G6 and G6*) | -71.3 ± 2.5 | 169.77 ± 1.68 | 64.07 ± 0.03 | 79.18 ± 4.02 |

REFERENCE

1. Hammond, N. B., B. S. Tolbert, R. Kierzek, D. H. Turner, and S. D. Kennedy. 2010. RNA internal loops with tandem AG pairs: the structure of the 5'GAGU/3'UGAG loop can be dramatically different from others, including 5'AAGU/3'UGAA. *Biochemistry*. 49(27):5817-5827, doi: 10.1021/bi100332r, http://www.ncbi.nlm.nih.gov/pubmed/20481618.
